# Congenital T cell activation impairs transitional to follicular B cell maturation in humans

**DOI:** 10.1101/2024.02.08.579495

**Authors:** Hugues Allard-Chamard, Kirsty Hillier, Michelle L. Ramseier, Alice Bertocchi, Naoki Kaneko, Katherine Premo, Tiffany Lam, Grace Yuen, Marshall Karpel, Vinay S. Mahajan, Christina Tsekeri, Jean Vencic, Rory Crotty, Anish Sharda, Sara Barmettler, Emma Westermann-Clark, Jolan E. Walter, Musie Ghebremichael, Alex K. Shalek, Jocelyn R. Farmer, Shiv Pillai

**Author notes:** Present address (Kirsty Hillier, Jocelyn R. Farmer). Authors contributed equally to this work (Hugues Allard-Chamard, Kirsty Hillier, Jocelyn R. Farmer, Shiv Pillai).

## Abstract

CTLA4-deficient patients exhibit profound humoral immune dysfunction, yet the basis for the B cell defect is not known. We observed a marked reduction in transitional to follicular B cell development in CTLA4-deficient patients, correlating with decreased CTLA4 function in regulatory T cells and increased mTORC1 signaling in transitional B cells. Treatment of transitional B cells with CD40L was sufficient to induce mTORC1 signaling and inhibit follicular B cell maturation *in vitro*. Frequent cell-cell contacts between CD40L^+^ T cells and naïve IgD^+^CD27^−^ B cells were observed in patient lymph nodes. Follicular B cell maturation in CTLA-deficient patients was partially rescued following CTLA4 replacement therapy *in vivo*. We conclude that functional regulatory T cells and the containment of excessive T cell activation are required for follicular B cells to mature and attain metabolic quiescence and thus acquire a state of immunological self-tolerance.

**One Sentence Summary:** Congenital T cell activation in CTLA4-deficient patients impairs transitional to follicular B cell maturation and can be rescued by CTLA4 replacement therapy *in vivo*.

## INTRODUCTION

Immature B cells emerge from the bone marrow as transitional B cells. A proportion of strongly self-reactive transitional B cells are clonally deleted, and the rest, widely assumed to be weakly self-reactive, differentiate into mature follicular B cells [1]. Mature follicular B cells recirculate, form the naïve B cell repertoire, and can subsequently be activated by CD4^+^ T cells during T-dependent immune responses to foreign antigens. Our studies on human transitional and follicular B cells have demonstrated that the developmental switch from the transitional to mature follicular B cell state is accompanied by the induction of the extracellular adenosine salvage pathway enzymes CD39 and CD73 and the activation of AMPK [2]. The consequent attenuation of mTORC1 activity leads to the acquisition of profound metabolic quiescence by follicular B cells [2]. Using a cohort of patients with hyperactivating mutations in *PIK3CD* causing activated PI3K-δ syndrome (ADPS), we have shown that constitutive mTORC1 signaling in human transitional B cells is sufficient to arrest development prior to the follicular B cell stage [2]. Whether extrinsic signals also influence transitional to follicular B cell development in humans or rodents is not known.

Cytotoxic T-lymphocyte-associated protein 4 (CTLA4) is a critical negative regulator of T cell co-stimulation that is constitutively expressed on T regulatory cells (Tregs). CTLA4 functions by competing with CD28 for surface CD80 and CD86 on antigen-presenting cells and removing these CD28 ligands through the process of trans-endocytosis [3]. CTLA4 can also act in cis, where upon sustained activation, effector T cells acquire surface CTLA4, which can restrain further TCR-dependent T cell activation [4, 5]. CTLA4 deficiency was first described in 2014 as an inherited, autosomal dominant condition of congenital Treg dysfunction causing constitutive T cell hyperactivation [6, 7]. Although *CTLA4* expression is restricted to T cells in humans, patients with CTLA4 deficiency present clinically with both humoral and cellular immune dysfunction. Defective humoral immunity is characterized by a decrease in circulating B cells, hypogammaglobulinemia, and recurrent infections. Pathological changes that also involve altered B cell function include robust autoimmune and lympho-infiltrative end-organ pathology (e.g. autoimmune cytopenias, entero-colitis, granulomatous and/or interstitial lung disease, dermatitis, and arthritis) [8]. Variable penetrance has been noted in CTLA4 deficiency [8], but T cell hyperactivation is observed even in asymptomatic carriers [9]. There is no evidence for the expression of *CTLA4* in nonmalignant human B lineage cells, including CTLA4-deficient patient B lineage cells [6], and interestingly mice with the engineered homozygous deletion of *ctla4* have no reported attenuation of B cell numbers or function.

The mechanism of T-cell-mediated autoimmune disease in CTLA4 deficiency stems logically from the direct or indirect role of CTLA4 in negatively regulating T cell activation and maintaining T cell tolerance in the periphery [10, 11]. In contrast, the mechanism of B cell dysfunction in CTLA4 deficiency remains poorly understood. Loss of total circulating B cells in CTLA4-deficient patients correlates with tissue sequestration and increased apoptosis of B cells, which express abnormally high levels of CD95, a marker of B cell activation [6, 7]. This suggests uncontrolled T-B collaboration and germinal center formation. Additionally, there is a loss of class-switched memory (IgD^−^CD27^+^) B cells and an expansion of CD21^lo^ B cells in circulation. Moreover, these CD21^lo^ B cells of CTLA4-deficient patients are CD38^lo^, distinguishing them from classical early transitional type 1 and 2 (T1/2) B cells, which are CD38^hi^ [7]. The exact ontology and contribution to disease pathogenesis of these CD21^lo^CD38^lo^ B cells is not known.

We show here that functional regulatory T cells are required by maturing B cells for the acquisition of metabolic quiescence at the mature follicular B cell stage.

## RESULTS

### Increased frequencies of transitional, activated Naive (aN), and marginal zone precursor (MZP) B cells in CTLA4-deficient patients

A cohort of patients with CTLA4 deficiency was longitudinally followed for development of low functional antibody levels in addition to autoimmune and end-organ lympho-infiltrative pathology, consistent with the disease spectrum of CTLA4 deficiency (Supplementary Table 1) [6–8, 12]. Symptomatic and asymptomatic carrier family members were additionally recruited. We performed peripheral blood B cell immune phenotyping by 16-parameter flow cytometry on our CTLA4-deficient patient cohort (Fig. 1) using the outlined panel and gating strategy (Supplementary Table 2 and Supplementary Fig. 1a). Consistent with previous reports describing CTLA4 deficiency, we observed a lower frequency of class-switched memory (IgD^−^CD27^+^) B cells and a corresponding higher frequency of B cells within the naïve gate (IgD^+^CD27^−^) in circulation (Fig. 1a, c) [6, 7]. Dissecting the naïve B cell compartment in greater detail, we observed increased transitional B cells in CTLA4-deficient patients, including T1/2 (CD10^+^MitoTracker^+^), T3a (CD10^−^MitoTracker^+^CD45RB^−^CD73^−^), and T3b (CD10^−^ MitoTracker^+^CD45RB^−^CD73^+^) B cells as we have recently defined them [2, 13]. In contrast, we found a significant reduction in MitoTracker^−^ follicular B cells in circulation (Fig. 1b, d), establishing an impairment of B cell maturation between the late transitional and follicular B cell stages. While circulating follicular B cells were lower in CTLA4-deficient patients, we observed higher frequencies of circulating activated naïve (aN; CD21^lo^CD11c^hi^) and marginal zone precursor (MZP; CD21^hi^CD45RB^+^) B cells within the naïve (IgD^+^CD27^−^) B cell compartment (Fig. 1b, d), consistent with the observation of extra-germinal center T-dependent B cell activation in CTLA4-deficient mice [14].

**Fig. 1.**
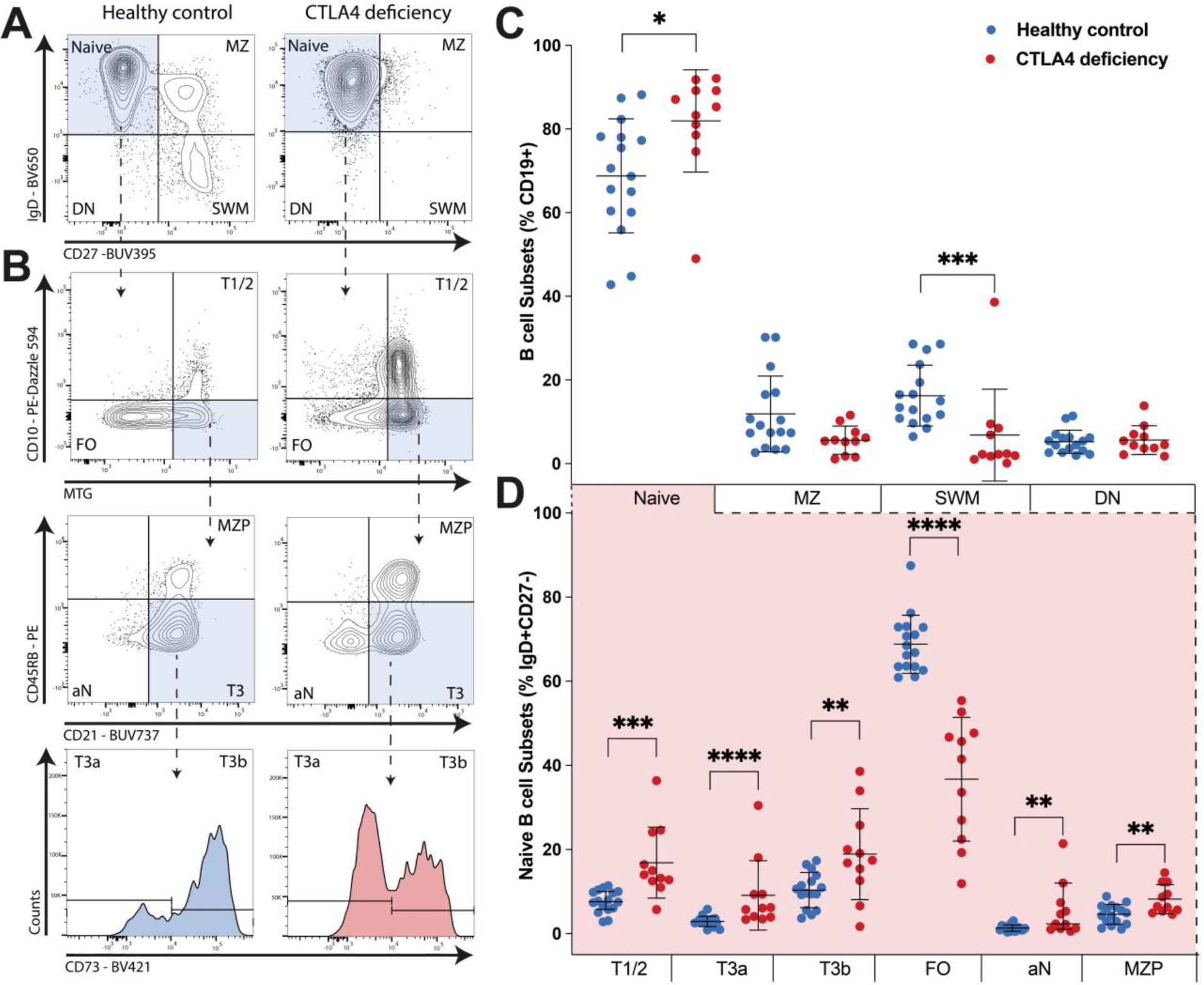
Increased frequency of circulating transitional B cells and a decrease in the frequency of circulating follicular B cells in CTLA4 deficiency. Flow cytometric analysis of peripheral blood B cell populations in CTLA4-deficient patients (n=11) and healthy controls (n=16). (A and B) Representative plots and gating strategy. (C) Quantification of major B cell subsets in peripheral blood, including Naïve, marginal zone (MZ), switched memory (SWM), and double negative (DN) populations as subset from total CD19^+^ B cells by the markers IgD and CD27. (D) Quantification of Naïve (IgD^+^CD27^−^) B cell subsets in peripheral blood, including T1/2, T3a, and T3b transitional B cells, resting mature follicular (FO) B cells, activated Naïve (aN) B cells, and presumed marginal zone precursor (MZP) B cells as further subsets by the indicated markers. Contour plots and histograms are representative of 11 CTLA4-deficient patients and 16 healthy controls. Symbols represent unique individuals; bars represent means (+/− SD) of all data. \**P* < 0.05; \*\**P* < 0.01; \*\*\**P* < 0.001; \*\*\*\**P* < 0.0001 by two-tailed Mann-Whitney U test.

### Circulating follicular B cell deficiency correlates with reduced Treg CTLA4 levels and impaired Treg function

Apart from the rare detection of CTLA4 in human B cells in the context of CD5^+^ monoclonal B cell malignancies [15, 16], *CTLA4* is understood to be primarily expressed by T cells. In our studies, intracellular flow cytometry revealed no discernible CTLA4 protein levels in transitional and follicular human B cells (MFI for intracellular CTLA4 levels <10^3^ in all naïve B cell subsets analyzed, Supplementary Fig. 2). We therefore hypothesized that the observed impairment in transitional to follicular B cell maturation was secondary to the congenital Treg dysfunction and resulting CD4^+^ T cell hyperactivation in CTLA4-deficient patients.

We tested the above hypothesis through the analysis of mutation-specific effects on Tregs in CTLA4-deficient patients as compared to healthy controls. Given the potential of heterozygous mutations in *CTLA4* to elicit a dominant negative effect, we additionally probed CTLA4 function in Tregs through analysis of CD80 trans-endocytosis [6]. All unique missense and non-coding region *CTLA4* mutations were observed to cause decreased CTLA4 levels and/or the attenuation of CTLA4-mediated trans-endocytosis in patient FOXP3^+^ Tregs as compared to healthy controls (Fig. 2). We further identified significant correlations between the loss of circulating follicular B cells and mutation-specific decreases in CTLA4 levels and CTLA4-mediated trans-endocytosis of CD80 (Fig. 2). For example, patients with the lowest CTLA4 levels in Tregs (homozygous *CTLA4* 3’ UTR mutation carriers) were observed to have the lowest circulating follicular B cell counts. Together, these data suggested that the loss of follicular B cells and the consequent humoral immune phenotype were directly proportional to the underlying magnitude of Treg dysfunction in patients with CTLA4 deficiency.

**Fig. 2.**
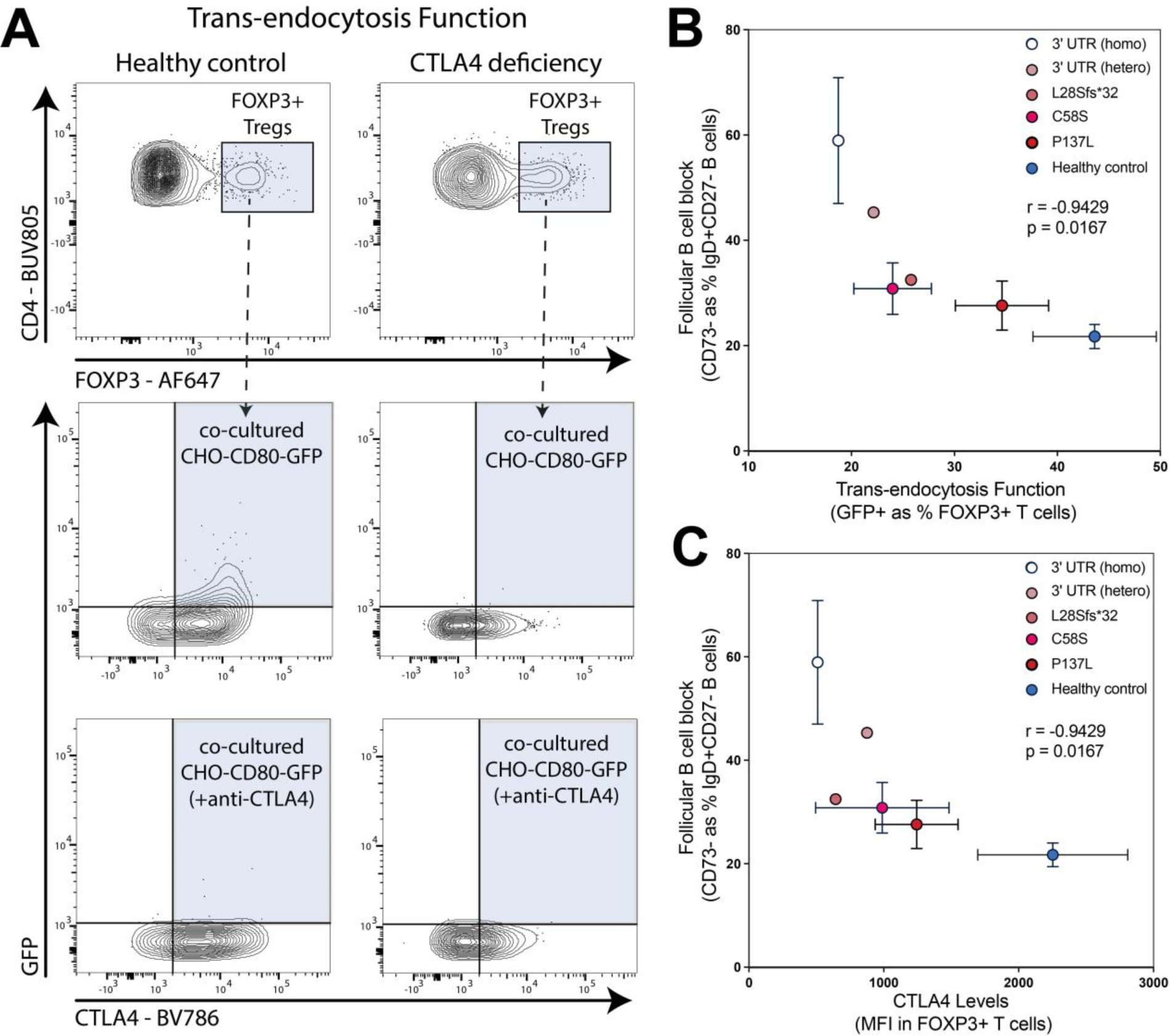
Reduced frequency of circulating follicular B cells correlates with degree of congenital Treg dysfunction in CTLA4 deficiency. CHO-CD80-GFP cells were co-cultured with activated T cells from the peripheral blood of 10 CTLA4-deficient patients and 14 healthy controls. Gated FOXP3^+^ Tregs that had acquired GFP from CHO expressing CD80-GFP cells were scored positive for intact CTLA4-mediated trans-endocytosis. (A) Representative contour plots and gating strategy. (B) Quantification of CTLA4 function by CD80-GFP trans-endocytosis (x-axis) and follicular B cell frequency in peripheral blood (data from Fig. 1; y-axis). Data were scored per unique individual and are grouped by underlying CTLA4 mutation type as compared to healthy controls. (C) Quantification of CTLA4 levels by intracellular staining in peripheral blood CD4^+^FOXP3^+^ Tregs (x-axis) and follicular B cell frequency in peripheral blood (data from Fig. 1; y-axis). Data were scored per unique individual and are grouped by underlying CTLA4 mutation type as compared to healthy controls. Contour plots are representative of 10 CTLA4-deficient patients and 14 healthy controls. Symbols represent mean value (+/−SD) per CTLA4 mutation type, as indicated, compared to healthy control. Correlation by nonparametric Spearman’s rank with correlation coefficient (r) and *P*-value shown.

### T cell hyperactivation and high CD40L levels in CTLA-deficient patients

We probed changes that occur in the T cell compartment of CTLA4-deficient patients using a combination of intra- and extra-cellular multiparameter flow cytometry (Supplementary Tables 3 and 4, gating strategy in Supplementary Fig. 1b). As is known in CTLA4 deficiency [6, 7], we observed loss of naïve CD4^+^ T cells and expansions of effector memory (EM), T effector memory CD45RA^+^ (TEMRA), and T follicular helper (TFH) CD4^+^ cells in circulation (Fig. 3a). Also consistent with what is known in CTLA4 deficiency, we observed an expansion of Tregs in circulation using both extracellular markers (CD25^hi^CD127^lo^) and the intracellular marker FOXP3 to identify CD4^+^ Tregs (Fig. 3b). Within the FOXP3^+^ Treg population, there was an expansion of CD25^lo^ Tregs in CTLA4-deficient patients compared to healthy controls (Fig. 3c) [7]. Probing for a known ligand that can induce B cell activation, we identified increased CD40L levels in effector CD4^+^ T cells (defined as CD4^+^CD56^−^CD45RA^−^) from CTLA4-deficient patients compared to healthy controls (Fig. 3d). Further T cell phenotyping also revealed NKT-like (CD3^+^CD56^+^) cell expansions in patients with CTLA4 deficiency (Supplementary Fig. 3).

**Fig. 3.**
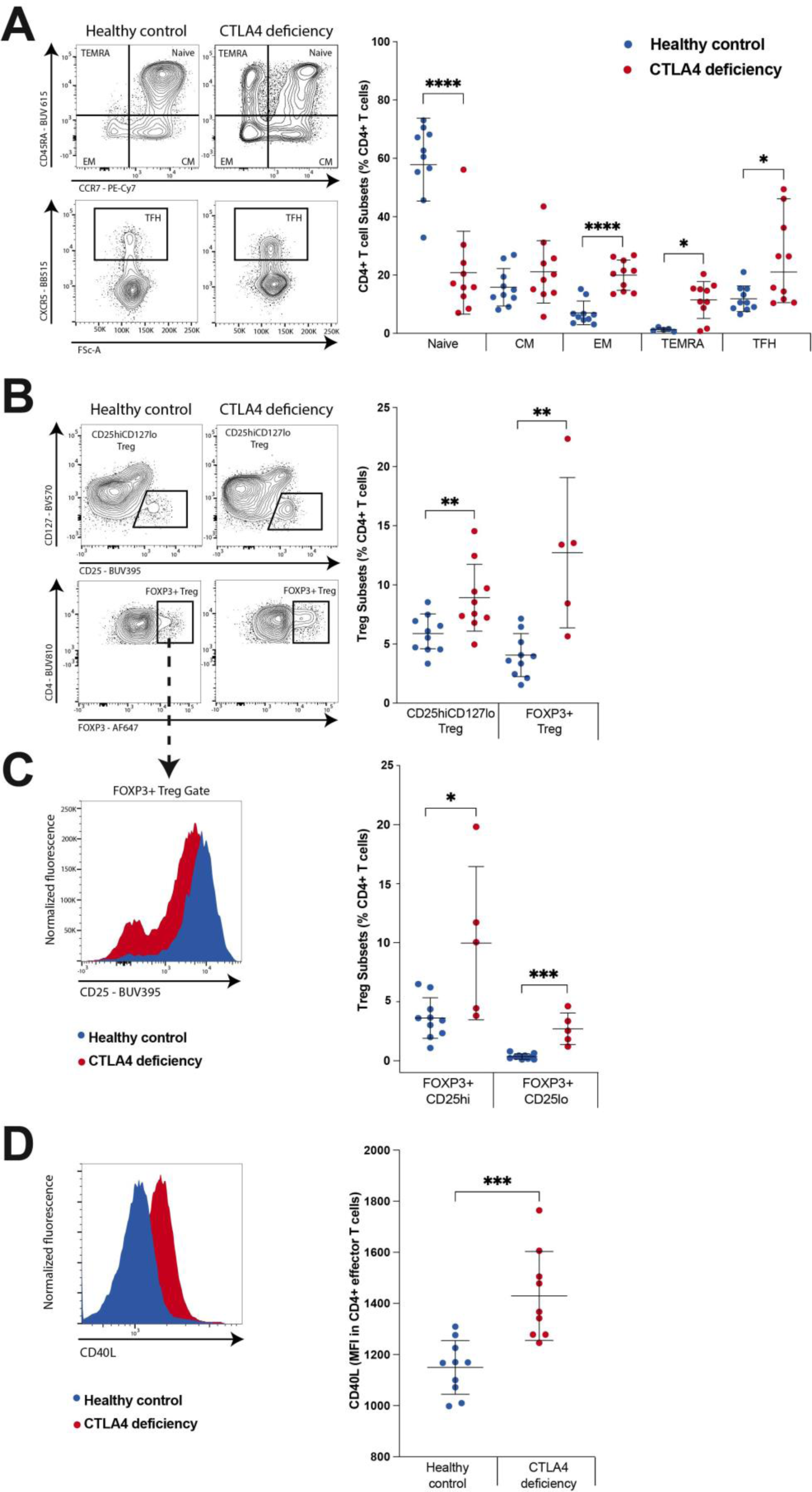
CTLA4 deficiency is associated with T cell hyperactivation and increased CD40L levels on activated CD4^+^ T cells. Flow cytometric analysis of peripheral blood T cell populations in CTLA4-deficient patients and healthy controls. (A) Analysis and quantification of major CD4^+^ T cell subsets, including naïve, central memory (CM), effector memory (EM), T effector memory CD45RA^+^ (TEMRA), and T follicular helper (TFH) cell populations as subsets from total CD4^+^ T cells by markers CD45RA, CCR7 and CXCR5. (B and C) Analysis and quantification of T regulatory cells (Tregs), which were CXCR5^−^ (excluding T follicular regulatory cells) using cell surface CD25 and CD127 and intracellular FOXP3 as markers (Supplementary Fig. 1b). (D) Analysis of CD40L levels in CD4^+^ T effector cells (CD4^+^CD56^−^CD45RA^−^), compared between CTLA4-deficient patients and healthy controls. Extracellular contour plots and histograms are representative of 10 CTLA4 patients and 10 healthy controls. Intracellular contour plots and histograms are representative of 5 CTLA4 patients and 10 healthy controls. Symbols represent unique individuals; bars represent means (+/− SD) of all data. \**P* < 0.05; \*\**P* < 0.01; \*\*\**P* < 0.001; \*\*\*\**P* < 0.0001 by two-tailed Mann-Whitney U test.

### Increased mTORC1 signaling defines aberrant B cell populations in CTLA4-deficient patients

Increased mTORC1 signaling is transcriptionally a hallmark of transitional B cell identity. Previous bulk RNA-sequencing (RNA-seq) analysis comparing flow-sorted circulating transitional B cells to resting follicular B cells revealed upregulation of B cell activation markers in the transitional cells, including *CD38*, *BCL2A1*, and *CD86*, and decreased expression of canonical follicular marker genes, such as *CCR7*, *FCER2,* and *IL13RA1* [2]. Here, follicular B cells were additionally defined by increased expression of *NT5E*, encoding for CD73, and *ABCB1* (Fig.4a). Both of these genes play critical roles in mediating ATP metabolism. Differentially expressed genes in the follicular B cell populations were enriched for NAD (nicotinamide adenine dinucleotide) signaling and biosynthesis pathways, in contrast to the strong enrichment of transitional B cell differentially expressed genes for mTORC1 signaling and oxidative phosphorylation pathways (Fig.4b). These results confirmed that maturation stage-restricted metabolic dependencies define transcriptional signatures separating transitional and follicular B cell populations [2].

We previously demonstrated that constitutive mTORC1 signaling in human transitional B cells is sufficient to impair development prior to the follicular B cell stage in a cohort of patients with gain-of-function mutations in PI3K-δ [2]. We therefore hypothesized that impaired transitional to follicular B cell maturation in CTLA4 deficiency could be driven by aberrant mTORC1 signaling in B cells downstream of CD4^+^ T cell activation. To resolve the dominant transitional B cell populations in CTLA4 deficiency and their underlying signaling patterns in transcriptional space, we performed single-cell RNA-sequencing (scRNA-seq) on the total naïve (IgD^+^CD27^−^) B cell compartment from four patients with CTLA4 deficiency and three healthy controls. Along with *CCR7*+ follicular (FO) B cells, clustering analysis resolved a subpopulation of transitional B cells (TrB), which most closely resemble the T3 transitional B cell stage given increased expression of *SOX4*, *CD38* and *HMCES*, in addition to sparse detection of the T1/2 marker *MME*. We additionally characterized a subpopulation of *ITGAX*^+^*TBX21*^+^ B cells. Although CD11c^hi^Tbet^+^ cells have sometimes been generically referred to as age-associated B cells (ABCs) and have been described during type I pathogen challenge and in autoimmunity (Fig.4c-e) [17, 18], it is clear that there are at least three distinct categories of *ITGAX^+^TBX21^+^* B cells in humans. In the IgD^+^CD27^−^ compartment these cells are described as activated Naïve (aN) B cells [19]. The CD27^+^*TBX21^+^ITGAX^+^*population exhibits a high level of somatic hypermutation and has been suggested by Wilson and colleagues to be germinal center derived [20]. In the IgD^−^CD27^−^ double negative compartment these ABC-like B cells are referred to as DN2 B cells [21]. Compared to healthy controls, decreases in the relative proportions of FO B cells in CTLA4-deficient patients were accompanied by increased proportions of both TrB cells and ABC-like aN B cells (Fig.4f-g), further corroborating our findings from flow analysis in Figure 1. The enrichment in T3 transitional B cells relative to T1/2 transitional B cells by scRNA-seq, as compared to flow cytometry – which relies on CD10 surface levels to largely define T1/2 transitional B cells – is consistent with known differences in B cell ontology as defined using transcriptome as compared to surface markers and reinforces using a combination technique to precisely define stages of B cell maturation. Using both approaches, we found that B cell maturation was impaired, specifically between late transitional and follicular B cell stages, in CTLA4-deficient patients as compared to healthy controls.

We next asked whether the ABC-like aN B cell population emerges in relation to transitional B cell populations, or whether these cells represent an independent subpopulation of B cells. Interestingly, ABC-like aN B cells shared expression of canonical transitional B cell marker genes, including *PLD4*, *MZB1*, and *CD1C* [22, 23]; furthermore, many genes that were most highly expressed in ABC-like aN B cells, such as *TNFRSF1B* and *CSGALNACT1* [24] were expressed to a lower extent in the TrB population and not expressed in follicular B cells (Fig.4d-e). We curated a signature of common differentially expressed genes in both TrB cells and ABC-like aN B cells compared to FO B cells (Fig.4h), many of which were also differentially expressed in bulk sorted transitional B cells compared to resting follicular B cells from healthy controls (Fig.4a). Scoring all cells by this signature similarly segregated both TrB cells and ABC-like aN B cells from their follicular counterparts in both healthy and CTLA4-deficient patients (Fig.4g), implicating that ABC-like aN B cells share core transitional B cell identity gene expression with the canonical TrB population (Fig.4i).

**Fig. 4.**
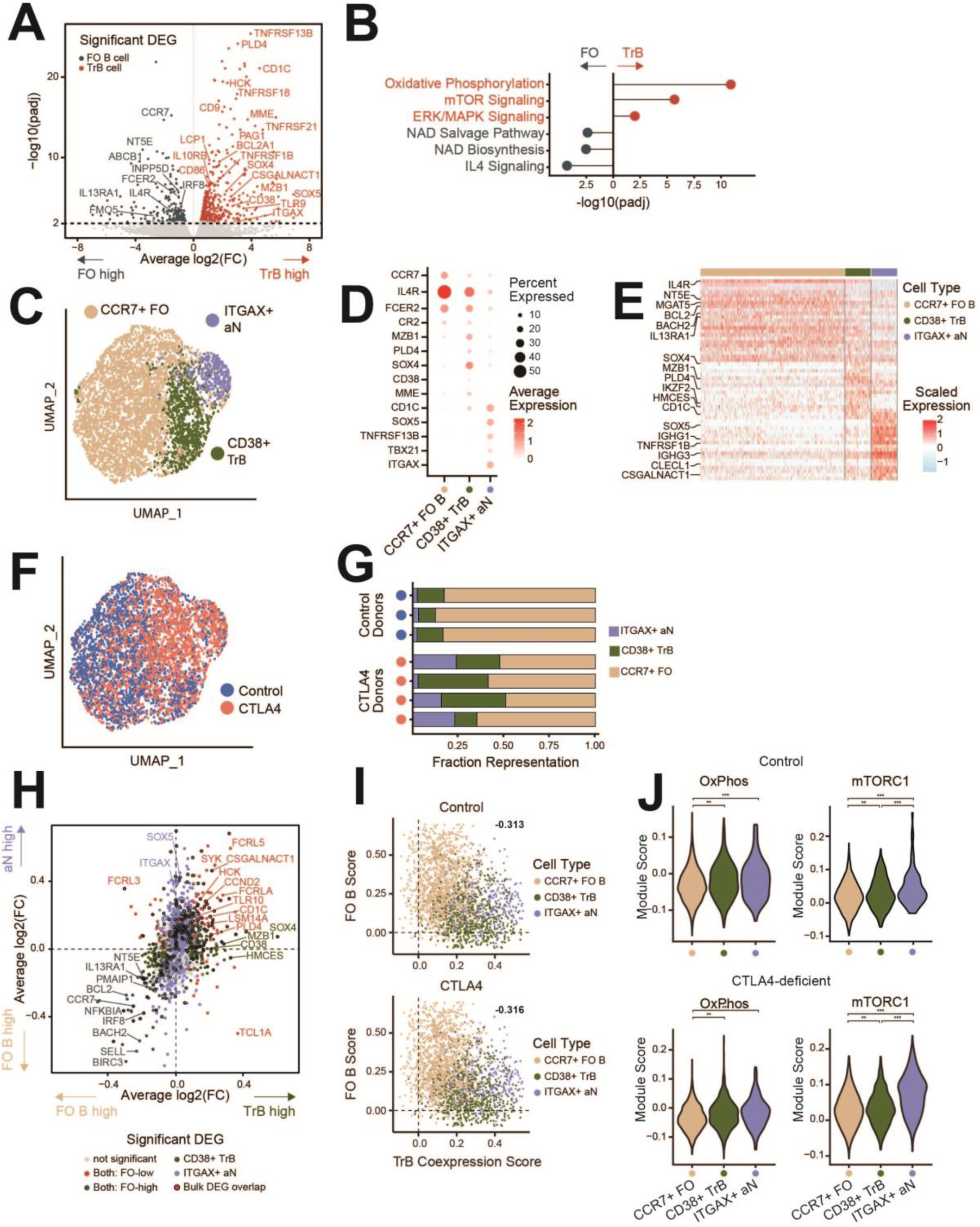
Increased mTORC1 signaling due to abundance of transitional and ABC-like aN B cell clusters in CTLA4-deficient patients. (A) Volcano plot of shared differential gene expression analysis between bulk RNA-seq samples of flow sorted resting follicular and transitional B cell populations (T1/2, T3) (n=3 healthy controls; significant differentially expressed genes defined by adjusted *P* value ≤ 0.01). (B) Adjusted *P* value of Ingenuity Pathway Analysis pathway overlaps with significantly differentially expressed genes in transitional B cells versus resting follicular B cells. (C) UMAP projection of total naïve (IgD^+^CD27^−^) B cells profiled by scRNA-seq from n=4 CTLA4-deficient patients and n=3 healthy controls, annotated by naïve B cell subtype. (D) Dot Plot showing the expression of follicular, transitional, and activated Naïve B cells by cell type annotation; color represents average normalized gene expression and size represents the percent cells with non-zero gene expression. (E) Heatmap of scaled gene expression of top 30 differentially expressed genes between annotated cell types. (F) UMAP projection of total naïve (IgD^+^CD27^−^) B cells profiled by scRNA-seq, annotated by control or CTLA4-deficient patients. (G) Fraction representation of each annotated cell type within each donor scRNA-seq sample. (H) Average log2-fold change of significant differentially expressed genes (adjusted *P* value ≤ 0.05) between *CD38*^+^ TrB cells versus *CCR7*^+^ FO B cells plotted on the x-axis, and average log2-fold change of significant differentially expressed genes between *ITGAX*^+^ ABC-like aN B cells versus *CCR7*^+^ FO B cells plotted on the y-axis, showing correlation in differential gene expression distinguishing both TrB cells and ABC-like aN B cells from FO B cells. Differentially expressed genes that overlap with significant differentially expressed genes from the bulk RNA-seq analysis in (A) are outlined. (I) Single-cell module scores defined by shared differentially expressed genes in both TrB and ABC-like aN cells plotted on the x-axis (TrB Coexpression score), versus module scores defined by shared differentially expressed genes in FO cells on the y-axis (FO B score). Single-cells from healthy controls are plotted separately from single-cells from CTLA4-deficient patients to demonstrate consistent trends in cell type population scores. Pearson correlation rho values (−0.313 for healthy controls, −0.316 for CTLA4-deficient patients) were both significant (*P* < 2E-16). (J) Violin Plots of module scores over msigDB Hallmark Oxidative Phosphorylation and mTORC1 Signaling pathways (see Methods). Violin Plots of single-cell module scores shown and evaluated for significance separately for healthy controls and CTLA4-deficient patients; \*\**P*<0.001, \*\*\**P*<0.0001.

Finally, given the relative accumulation of both TrB and ABC-like aN B cells in CTLA4-deficiency, we explored whether the ABC-like aN B cells utilized the same underlying metabolic signatures as transitional B cells. By module scoring our B cell subpopulations, we found increased transcriptomic signatures of oxidative phosphorylation in both TrB cells and ABC-like aN B cells compared to FO B cells. Furthermore, we observed elevated mTORC1 signaling in TrB cells compared to FO B cells, and further increased mTORC1 signaling in ABC-like aN B cells compared to both TrB and FO B cells. These trends were consistent in both healthy and CTLA4-deficient patient B cells, implicating that mTORC1 signaling strongly defines ABC-like aN B cells (Fig.4j). Collectively, these findings implicate a potential transitional B cell origin for ABC-like aN B cells and nominate mTORC1 signaling as both an inhibitory pathway for normal follicular B cell maturation and permissive for ABC-like aN B cell maturation in the context of increased CD4^+^ T cell activation.

### CD40L induces mTORC1 signaling and inhibits transitional to follicular B cell maturation *in vitro*

We hypothesized that, in the context of CTLA4 deficiency, mTORC1 hyperactivation may be occurring secondary to CD40 signaling on B cells engaged by activated CD40L^+^ T cells. Using an *in vitro* B cell differentiation model, we have previously shown that transitional to follicular B cell maturation is enhanced with AMPK stimulation and mTORC1 inhibition [2]. Using this model system, we now probed the direct effects of CD40L on transitional B cell signaling and developmental potential. We observed increased levels of phosphorylated S6 in response to CD40L treatment of transitional B cells (Fig. 5a), which suggested that CD40L is sufficient to induce mTORC1 signaling in transitional B cells. Using our *in vitro* assay of transitional to follicular B cell maturation [2], we additionally probed developmental potential and observed decreased generation of follicular B cells in transitional B cell cultures treated for 24 hours with CD40L compared to those left untreated (Fig. 5b). Together with previously published data, these data suggested that, while mTORC1 inhibition augments transitional to follicular B cell maturation *in vitro* [2], induction of phosphorylated S6 signaling by CD40L in transitional B cells drives the opposite phenotype and limits their potential to mature to follicular B cells *in vitro*.

**Fig. 5.**
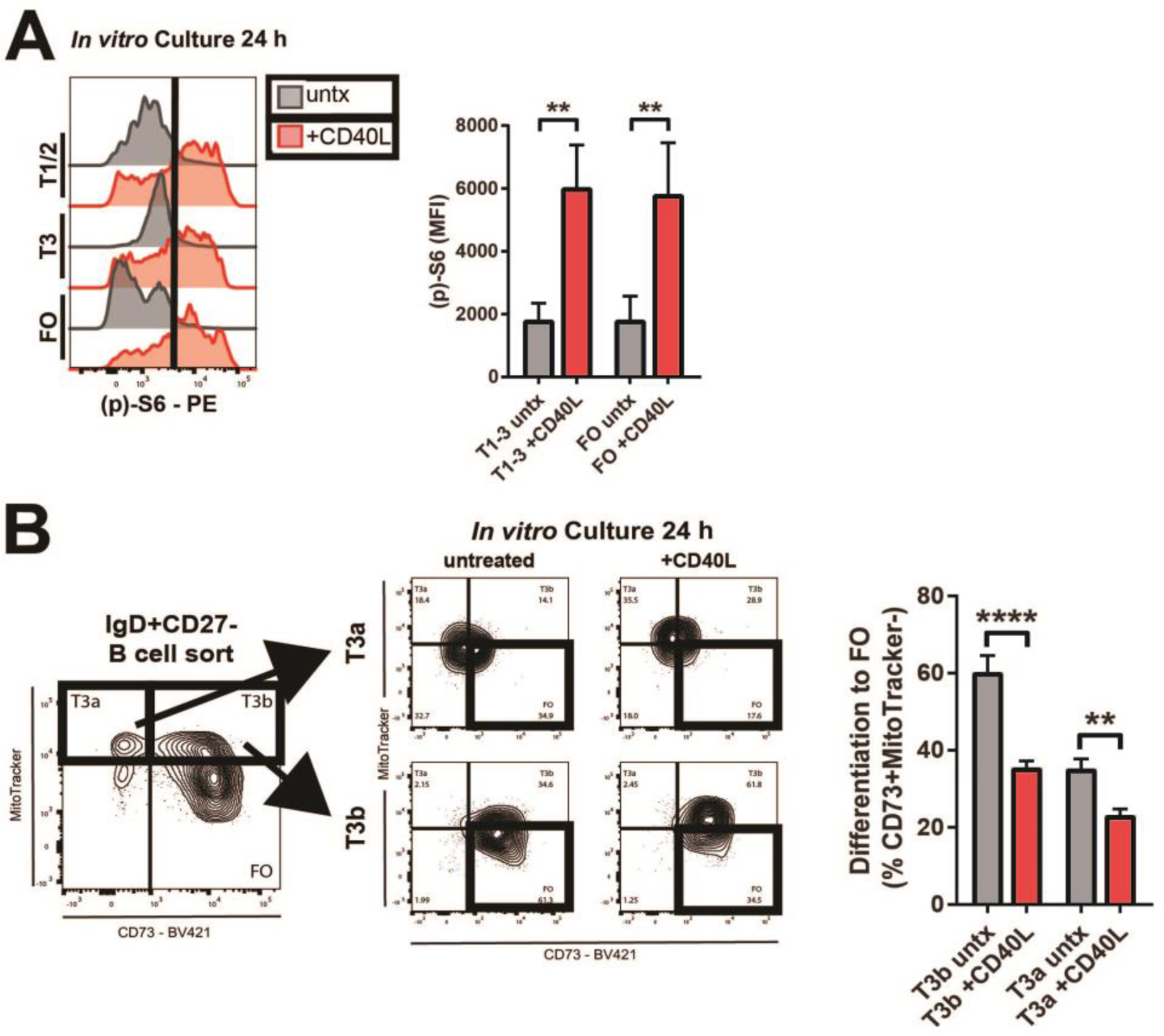
CD40L+ T cells can contact transitional B cells, induce mTORC1 signaling, and arrest follicular B cell maturation. (A) Levels of phosphorylated (p)-S6 by flow cytometry in transitional (T) and follicular (FO) B cell subsets, sorted and cultured for 24 hours *in vitro* with compared to without CD40L treatment as indicated. Data are representative of three independent experiments. Quantified data are means (+/− SD) of all data. (B) Differentiation of T3a and T3b B cells, sorted and cultured for 24 hours *in vitro* with compared to without CD40L treatment as indicated. Differentiation to the FO B cell stage by flow cytometry was scored as shown by the acquisition of the CD73^+^MTG^−^ phenotype. Data are representative of three independent experiments. Quantified data are means (+/− SD) of all data. \**P* < 0.05; \*\**P* < 0.01; \*\*\*\**P* < 0.0001 by unpaired Student’s *t* test.

We probed where an interaction of CD40L^+^ T cells and naïve B cells could be occurring *in vivo*. We examined lymph nodes of patients with common variable immunodeficiency (CVID) caused by a congenital syndrome of T cell activation (APDS and CTLA4 deficiency). We observed cell-cell contacts between CD40L^+^ T cells and naïve (IgD^+^CD27^−^) B cells occurring prominently in the T cell zones that were far more frequent in patients with congenital T cell activation as compared to secondary lymphoid organs of healthy controls (Supplementary Fig. 7). These data suggested that activated T cells can directly interact with naïve (IgD^+^CD27^−^) B cells in human lymphoid organs.

### Follicular B cell maturation is restored with CTLA4-Ig treatment

In a subset of our CTLA4 cohort, onset of autoimmune and lympho-infiltrative disease necessitated the start of abatacept (CTLA4-Ig) therapy. Robust clinical response to abatacept has been demonstrated in CTLA4 deficiency [8, 25], as well as in disorders that phenocopy this condition, homozygous LRBA and DEF6 deficiencies [26–28]. Clinical response to abatacept in our cohort was significant, including amelioration of autoimmune cytopenias, entero-colitis, arthritis, lymphadenopathy, and granulomatous-lymphocytic interstitial lung disease (GLILD). Dose and duration of treatment are detailed in Supplementary Table 1. In patients treated for at least three months with systemic abatacept therapy, we observed a robust decrease in circulating early transitional (T1/2) B cells and a corresponding increase in follicular B cells. This correction in B cell maturation restored follicular B cell counts in CTLA4-deficient patients to levels seen in healthy control subjects (Fig. 6a).

**Fig. 6.**
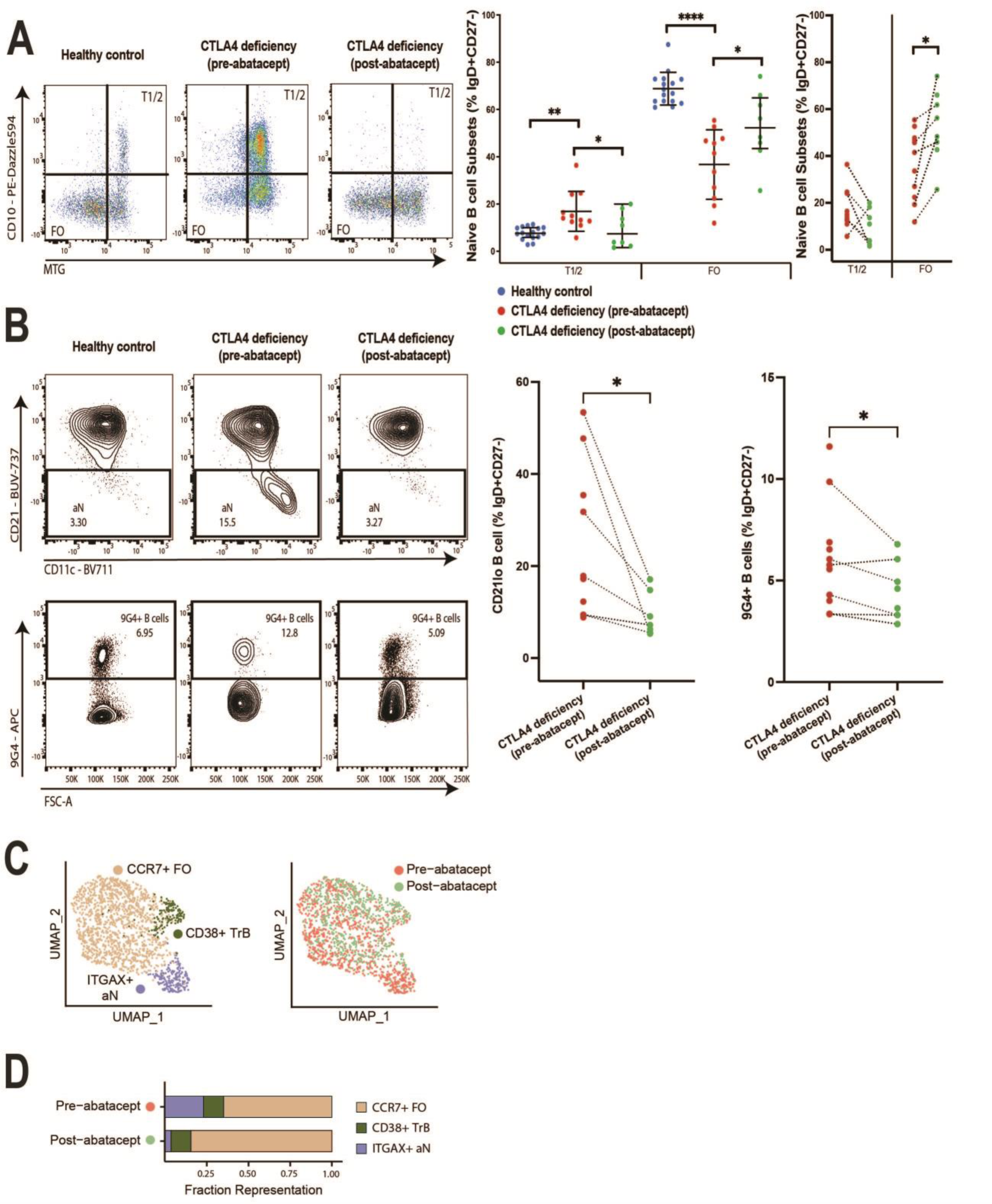
Treatment with abatacept (CTLA4-Ig) restores follicular B cell maturation coincident with decreased CD40L levels on effector T cells in CTLA4 deficiency. (A, B) Flow cytometry analysis of B cell populations before and after abatacept treatment in CTLA4 deficiency. B cell plots are representative of 11 patients prior to treatment and 8 treated patients (6 captured both pre- and post-therapy) compared to 16 healthy controls. (C) Single-cell RNA sequencing (scRNA-seq) of total IgD^+^CD27^−^ B cells. UMAP projection of cells is shown by cell type clustering and state of abatacept treatment. (D) Fraction representation of each clustered scRNA-seq cell type. Quantified data are means +/− SD of all data. \**P* < 0.05; \*\**P* < 0.01; \*\*\**P* < 0.001; \*\*\*\**P* < 0.0001 by ANOVA for multiple comparisons and paired Wilcoxon rank sum test for pair-wise comparisons, respectively.

Coincident with the restoration of follicular B cell counts in circulation, we observed decreased generation of activated and autoreactive B cell subsets following abatacept therapy in our CTLA4-deficient patients (Fig. 6b). After abatacept therapy, we observed lower frequencies of CD21^lo^ B cells and fewer autoreactive naïve B cells, as defined by binding to the 9G4 substrate, within the IgD^+^CD27^−^ naïve B cell gate (that includes the activated naïve B cell population) in the blood of our CTLA4-deficient patients. Accounting for high inter-patient variability, we performed scRNA-seq on the naïve B cell compartment of individual treated subjects and compared pre- to post-therapy naïve (IgD^+^CD27^−^) B cell transcriptomes. Using unbiased UMAP clustering, a distinct B cell subpopulation was identified pre-treatment with abatacept. This distinct pre-treatment B cell subpopulation was observed to be CD11c^hi^CD21^lo^T-bet^+^, and contained transcripts from IgA and IgG loci, suggesting the initiation of class switching. The abatacept-mediated change in B cell differentiation was coincident with resolution of the high CD40L levels previously observed on effector T cells to those seen in healthy controls (Fig. 6c-d). In contrast to the robust phenotypic change observed in the B cell compartment, total populations of EM and TEMRA CD4^+^ T cells remained significantly unchanged pre- to post-treatment with abatacept (Supplementary Fig.4). Together, these data suggested baseline extra-follicular T-dependent B cell activation in CTLA4 deficiency and the restoration of normal follicular B cell differentiation following three or more months of systemic abatacept therapy.

## DISCUSSION

These studies have helped to reveal how CD4^+^ T cell effectors in conjunction with Tregs can regulate a specific checkpoint during human B cell development. This work also provides some insights regarding the contribution of Tregs to the maintenance of peripheral B cell tolerance at the transitional B cell stage, indicating that in human CTLA4 deficiency there is a previously unrecognized impairment in early B cell development that likely is a major contributor to the poorly explained loss of humoral immunity seen in these patients. This work also highlights the importance of refining markers to discriminate between transitional and follicular B cells. CD24 and CD38 have been widely used in flow cytometry to discriminate transitional from follicular B cells but are not able to separate late transitional (T3) from follicular B cells, specifically [2]. The additional use of MitoTracker and CD10 allows for the further resolution of late T3 transitional and follicular B cells, the latter of which are MitoTracker^−^ due to the acquisition of the ABCB1 efflux transporter. CD10^−^MitoTracker^+^ T3 B cells can further be divided based on CD73 expression into T3a (CD73^−^) and T3b (CD73^+^) as we have defined them [2]. The acquisition of surface CD73 is of developmental significance, since in culture CD73^+^ transitional B cells spontaneously mature into follicular B cells while CD73^−^ transitional B cells do not [2].

CTLA4 deficiency is a known state of congenital Treg dysfunction with coincident chronic hyperactivation of effector T cells. Despite restricted *CTLA4* expression to the T cell lineage in humans, CTLA4 deficiency is associated with a well described humoral immune dysfunction, specifically an inability to generate memory B cell responses to both T-dependent and T-independent antigen challenges [6–8]. CTLA4-deficient patients are lymphopenic, with a reduction in class-switched memory B cells and the expansion of activated CD21^lo^ B cells in the circulation [6, 7]. While the molecular basis for this B cell phenotype is not known, possible mechanisms include increased B cell apoptosis [29], robust B cell trafficking to end-organs, and/or impaired B cell differentiation. We herein have demonstrated an increased frequency of circulating transitional B cells with a corresponding loss of follicular B cells in CTLA4-deficient patients. The most significant increase in comparison to healthy controls occurred at the T3a transitional B cell stage. As these transitional B cells are MitoTracker^+^CD10^−^, these data are consistent with the lack of any previous description of impaired transitional to follicular B cell development, since previous studies only used CD10, CD21, and CD38 surface levels to phenotype B cells in the naïve (IgD^+^CD27^−^) gate in CTLA4-deficient patients and to thus define any transitional B cells within this gate [7]. Our data support a model of dysregulated B cell differentiation, specifically between the late transitional and follicular stages, as an underlying basis for the decreased humoral immune protection observed in patients with CTLA4 deficiency. We cannot fully exclude the alternative hypothesis that mature follicular B cells are sequestered in tissue, inflating the relative proportion of transitional to follicular B cells found in the peripheral blood. However, as CTLA4-deficient patients are clinically defined by the loss of antigen-specific B cell responses, our data are well aligned with the state of functional follicular B cell compromise previously described in CTLA4 deficiency [6, 7]. Moreover, our data support the redirection of transitional B cells to an activated Naïve (aN) fate, which as well coincides with published data in CTLA4 deficiency and larger common variable immunodeficiency (CVID), demonstrating the role of CD21^lo^ B cells in tissue-infiltrative end-organ disease [30].

Various autoimmune clinical manifestations are seen in CTLA4-deficient patients, including prominent infiltration of nonlymphoid organs, including the lung, liver, and gastrointestinal lumen, by expanded CD3^+^ T cell and CD19^+^ B cell populations [6–8]. Autoreactive B cells as well as overt autoantibody-mediated disease sequelae such as autoimmune cytopenias and autoimmune thyroiditis additionally are known to occur in CTLA4-deficient patients [31]. Given that B cell maturation is impaired prior to the follicular stage, one hypothesis for how CTLA4-deficient patients progress to B cell-mediated autoimmunity is that while circulating follicular B cells were lower in CTLA4-deficient patients, we observed higher frequencies of circulating activated Naïve (aN) B cells, consistent with the observation of extra-germinal center B cell activation in CTLA4-deficient mice [14]. These data are aligned with the description of extra-follicular T-dependent B cell differentiation in other systemic autoimmune conditions, such as systemic lupus erythematosus (SLE) [32], as well as in conditions of robust immune stimulation such as viral infection and immunization [33, 34]. Patients with partial RAG deficiency exhibit an expansion of CD27^−^IgD^hi^CD38^−^CD24^−^IgM^+^CD21^lo^ activated naïve B cells as well as Tbet^+^ age-related B cells (ABCs) that are detectable in the circulation, and in follicular and extrafollicular areas [17]. This alternative, extrafollicular B cell differentiation pathway can result in the production of class-switched, self-reactive antibodies capable of mediating end-organ autoimmune disease, as we also observe here in patients with CTLA4 deficiency. In our current studies, scRNA-seq revealed a cluster of *ITGAX^+^TBX21^+^*B cells with transcripts from IgA and IgG loci, suggesting the initiation of class switching, most consistent with ABC-like aN B cells. It is likely that these cells coincide with the cells identified as activated naïve B cells by flow cytometry (CD21^lo^CD11c^hi^) although intrinsic limitations of flow cytometry granularity limit this correlation. In single cell analysis, these ABC-like aN B cells also shared genes expressed with transitional B cells. The relationship between transitional B cells and aN B cells may be further supported by single cell trajectory analysis, and is a focus of ongoing work.

We have shown that intrinsic hyperactivation of the mTORC1 pathway can be driven by the constitutive activation of PI3K-δ [2]. Failure to attenuate mTORC1 signaling prevents B cells from completing their differentiation to the follicular B cell stage. Similar to PI3K-δ gain-of-function patients, CTLA4-deficient patients demonstrated a transcriptomic signature of increased mTORC1 activation in naïve (IgD^+^CD27^−^) B cells. Specifically, through scRNA-seq we observed increased transcriptomic signatures of oxidative phosphorylation in transitional B cells and ABC-like aN B cells compared to FO B cells, with elevated mTORC1 signaling in ABC-like aN B cells and transitional B cells compared to FO B cells. Moreover, CD40L was sufficient to both induce phosphorylated S6 signaling in transitional B cells and to impair their *in vitro* potential to form follicular B cells. These data support one model whereby transitional B cells are developmentally impaired in CTLA4-deficient patients through the interaction “in trans” with CD40L on activated effector CD4^+^ T cells, which may also facilitate the T-dependent differentiation in CTLA4-deficient patients and healthy controls of activated Naïve B cells. It might be speculated that these activated CD4^+^ effectors T cells represent unrestrained self-reactive CD4^+^ T cells that can provide cognate extra-follicular help to antigen-specific self-reactive B cells. It has been suggested that a large fraction of naïve B cells in humans are self-reactive and thus susceptible to such interaction [35, 36]. While we do note an expansion of identifiable self-reactive 9G4^+^ B cells, our studies do not address the antigen specificity of the expanded CD4^+^ T cells seen in the patients we have studied. Furthermore, we investigated expression of VH4-34 with the 9G4 reagent in a limited manner in this manuscript and more detailed analyses of B cell autoreactivity in the context of CTLA4 deficiency is the subject of ongoing work.

We also explored where one site of interaction between early peripheral B cells and CD40L^+^ T cells can occur *in vivo*. Late transitional B cells acquire CCR7 expression (Supplementary Fig. 5). In the lymph node, these cells initially follow a gradient of CCL19 and CCL21, cross high endothelial venules and are drawn first into outer paracortical area in the T cell zone, from where they are rapidly induced to enter the B cell follicle based on their expression of CXCR5. We have shown here that T cell-B cell-interaction events do occur between naïve B cells and CD40L-expressing T cells in the T cell zone of lymph nodes of CVID patients, including CTLA4-deficient patients. Additionally, these deficient patients have prominent infiltration of nonlymphoid organs with expanded T and B cell populations [6–8]. In the lung, specifically, these B cell infiltrates have been shown to organize into pseudo germinal centers – which are likely extrafollicular T cell-B cell aggregates. So even while promiscuous interaction of T cells with B cells prevents their developmental maturation in secondary lymphoid organs, T cell to B cell interactions also likely drive pathological events in the nonlymphoid end-organs of CTLA4-deficient patients. The different phenotypes seen in mice and humans could perhaps be also accounted for by species-specific differences in the circulation of B cells in lymphoid organs. In mice, the splenic marginal zone is better defined than in humans and anatomical differences in B cell niches between mice and human may also contribute to the different species-specific phenotypes of mutations that influence T-B collaboration [37].

Our results are biologically consistent with the data from Savage and colleagues indicating that in healthy mice and humans a large proportion of T follicular helper cells in lymph nodes are intrinsically self-reactive but are held in check by regulatory T cells [38]. These data are consistent with previous studies from Germain and colleagues that have shown that in lymph nodes there are local regulatory T cell feedback circuits that prune self-reactive T cells and maintain homeostasis [39]. It is this disordered homeostasis in CTLA4-deficient patients that likely permits the promiscuous activation of transitional B cells, as they enter the periarteriolar lymphatic sheaths, and the resulting impairment of normal differentiation that we have observed in our studies.

Limitations to this study include the lack of exact aged-matched healthy controls. While B cell subsets in the peripheral blood of healthy volunteers vary by age, most differences are seen in the very young and very old demographics [40]. Here, median age of both groups was in early adulthood (40.5 compared to 23 years for healthy controls versus CTLA4-deficient patients). Moreover, we were able to replicate multiple B cell flow phenotypes already published in CTLA4 deficiency, including the loss of class-switched memory B cells and the expansion of naïve B cells. Although herein we show that CD40L is sufficient to impair transitional to follicular B cell maturation *in vitro*, we accept that other ligands not evaluated in this study may have a contributory role in transitional to follicular B cell maturation. Studying B cell differentiation *in vitro* will never fully recapitulate the differentiation potential of human B cells *in vivo*, and there is a paucity of suitable model systems given the known differences between human and murine transitional B cell compartments noted above. Direct comparison of tissues and draining lymph nodes would be needed to confirm increased interactions between naïve B cells and CD40L+ T cells in CVID patients and is a goal of future work. Finally, CTLA4 is expressed constitutively on the surface of CD5^+^ B-1a cells in mouse [41]. While the existence of B-1 cells in the circulation of adult humans remains controversial, CTLA4 expression has been observed in CD5^+^ B cells of patients with chronic B-cell lymphocytic leukemia [15, 16]. It is therefore theoretically possible that inherited CTLA4 deficiency could cause direct transitional B cell dysfunction that is corrected by CTLA4-Ig replacement therapy. However, we failed to demonstrate discernable intracellular RNA levels of CTLA4 in naïve (IgD^+^CD27^−^) B cells, including transitional and follicular B cell subsets, specifically, and intracellular protein levels were not reliably detectable in T3/FO B cells by flow cytometry. Additionally, it is theoretically possible that CTLA4-Ig therapy mediates a direct effect on transitional B cells through the binding of B cell co-stimulatory molecules (CD80/CD86). CTLA4-Ig has been shown to alter CD80 and CD86 levels on *in vitro* cultured total CD19^+^ B cells as well as mediate intracellular signaling through this interaction [42, 43]. However, our analysis of surface CD86 levels on naïve B cell subsets demonstrated extremely low abundance on transitional and follicular B cells, which make this probability less likely (Supplementary Fig. 6). In contrast, we observed maturational impairment of transitional B cells on their way to a follicular B fate that correlated in a linear manner with lower CTLA4 levels and diminished CTLA4-dependent trans-endocytosis in FOXP3^+^ Tregs. Moreover, we observed direct cell-cell contacts between CD40L^+^ activated T cells and naïve (IgD^+^CD27^−^) B cells in the T cell zones of patient lymph nodes. These data suggest that functional regulatory T cells are required for the maturation of metabolically quiescent self-tolerant mature follicular B cells. We observed a decrease in T1/2 B cells and an increase in FO B cells, which was corrected with CTLA4 Ig (abatacept) therapy by flow cytometry. Additionally, we observed a decrease in transitional B cells and *ITGAX*^+^ ABC-like aN B cells post-abatacept therapy by scRNA sequencing analysis. These findings indicate that abatacept (CTLA4-Ig) therapy will likely correct the pathologic B cell activation observed in CTLA4-deficient patients.

## MATERIALS AND METHODS

### Recruitment of patients with CTLA4 mutations and mutation analysis

This research was performed in accordance with the Mass General Brigham Institutional Review Board (protocols 2011P000940 and 2018P001584). From primary immune deficiency cohorts and their family members followed in the outpatient clinical practices at the Massachusetts General Hospital [44] and the University of South Florida, patients identified as potential CTLA4 deficiency by next-generation panel or whole exome sequencing were approached for inclusion in the study. Written informed consent prior to specimen collection was obtained from all patient participants included in this study. CTLA deficiency was validated using intracellular CTLA4 flow cytometry analysis in combination with CD80 trans-endocytosis functional analysis in FOXP3^+^ Tregs as detailed (Fig. 2). Healthy adults without any history of recurrent infection or autoimmunity were recruited as controls. Median age of healthy controls was 40.5 years (IQR 33.25-45 years). Median age of CTLA4-deficient patients was 23 years (IQR 17-31 years). Exact numbers of CTLA4-deficient patients and healthy controls used per assay are detailed exactly in each figure legend and in Supplemental Table 1.

### B cell Profiling by Flow Cytometry in CTLA4 deficiency

The distinction between transitional and follicular human B cells was accomplished by ABCB1 transporter efflux profiling, using MitoTracker™ Green (MTG; ThermoFisher Scientific) [2, 45, 46], in combination with CD73 surface staining [2, 13]. In brief, 5 million previously frozen peripheral blood mononuclear cells (PBMCs) were incubated in RPMI 1640 media containing MTG at a 1:10,000 dilution (100-nM concentration) for 30 min at 37°C. Subsequently, cells were spun down at 1,250 rpm for 6 min and rested in fresh RPMI 1640 media for an additional 30 mins at 37°C to allow efflux of the dye in ABCB1^+^ cells. Cells were blocked for 15 min with 10 µL of human Fc receptor blocking solution (TruStain FcX^TM^; BioLegend). To minimize the interaction of polymeric dyes and improve discrete fluorochrome readout, Brilliant Stain Buffer Plus (BD Biosciences) was included at 10 µL per sample. Cells were surface-stained with optimized concentrations of fluorochrome-conjugated primary antibodies for 30 min at 4°C in DMEM no phenol red (21063029, ThermoFisher Scientific) supplemented with 2% fetal bovine serum (FBS). All washes were carried out using phosphate-buffered saline (PBS) supplemented with 0.5% bovine serum albumin (BSA). DRAQ7T^M^ (BioLegend) was used to exclude dead cells as per the manufacturer’s instructions. Compensation was performed using VersaComp Antibody Capture Beads (Beckman Coulter) apart from MTG, which required staining or not of Ramos cells [American Type Culture Collection (ATCC)], which are negative for the ABCB1 efflux pump. For analysis, B cells were run on a five-laser BD FACS Symphony (5S Symphony). B cells were sorted on a FACS Aria II (BD Biosciences) cell sorter and collected in RMPI 1640 media supplemented with 20% FBS. Primary B cell panel is listed in Supplementary Table 2. In a subset of experiments (Fig. S2), CTLA4 (BNI3; BD Biosciences; 1:20) was substituted into the panel using an additional intra-cellular staining step as detailed for T cells below. In a subset of experiments (Figs. S5, S6), CCR7 (G043H7; BioLegend, 1:10) and CD86 (FUN-1; BD Biosciences; 1:40) were substituted into the panel, respectively.

### T cell Profiling by Flow Cytometry in CTLA4 deficiency

For T cell staining, a similar protocol was applied, with few notable changes. For intra-cellular staining only, viability dye (Live/Dead Blue, Invitrogen) was added to exclude dead cells as per the manufacturer’s instructions. After Fc block, cells were stained for chemokine receptors with optimized concentrations of fluorochrome-conjugated primary antibodies for 30 min at 37°C. Cells were then washed with PBS 0.5% BSA, spun down at 1,250 rpm for 6 min, and further surface-stained with optimized concentrations of fluorochrome-conjugated primary antibodies for 30 min at 4°C. Following the extracellular staining protocol, we proceeded with intracellular staining in a subset of experiments. The cells were re-suspended in fixation and permeabilization buffers using the True-Nuclear^TM^ Transcription Factor Buffer Set (BioLegend) per manufacturer’s instructions. Cells were stained with fluorochrome-conjugated primary antibodies for intracellular staining for 30 min at room temperature. For analysis, T cells were run on a five-laser BD FACS Symphony (5S Symphony) analyzer. Primary T cell panels are listed in Supplementary Tables 3 and 4.

### CD80 Trans-endocytosis Assay

CHO cells stably overexpressing CD80-GFP were kindly provided by the Sansom laboratory [3]. CHO-CD80-GFP cells or CHO cells (ATCC® CCL-61) as control were plated to 75% confluence in 48-well dishes and stained immediately prior to T cell co-culture with CellTrace Violet (Cell Proliferation kit; ThermoFisher Scientific) according to manufacturer’s instructions. Primary human CD4^+^ T cells from CTLA4-deficient patients or healthy controls were purified from 5 million frozen PBMCs using a biotin labeled anti-human CD4 antibody (OKT4; BioLegend; 1:20) followed by magnetic separation with anti-biotin microbeads (Miltenyi Biotic) according to manufacturer’s instructions. Purified CD4^+^ T cells were co-cultured with CHO cells for 16 hours in the presence of T cell activation using anti-CD3/CD28 Dynabeads™ (ThermoFisher Scientific) at an estimated 1:1 bead-to-cell ratio. In a subset of the experiment, recombinant human CTLA4-Fc chimera fusion protein (R&D Systems) was added at a concentration of 5 ug/mL to inhibit CTLA4-mediated trans-endocytosis. Flow cytometry analysis of CTLA4-mediated trans-endocytosis was performed as previously described [6] using the True-Nuclear^TM^ Transcription Factor Buffer Set (BioLegend) and the following intracellular antibody panel: CD25 (2A3; BD Biosciences; 1:20), CD4 (SK3; BD Biosciences; 1:40), CXCR5 (J252D4; BioLegend; 1:30), CTLA4 (BNI3; BD Biosciences; 1:20), anti-GFP (FM264G; BioLegend; 1:200), CD127 (A019D5; BioLegend; 1:30), and FOXP3 (259D/C7; BD Biosciences; 1:20). FOXP3^+^ Tregs that had acquired GFP from CHO expressing CD80-GFP cells were scored positive for intact CTLA4-mediated trans-endocytosis.

### *In vitro* B cell differentiation and phospho–flow cytometry

PBMCs isolated from fresh healthy donor human leukopacks were surface labeled as previously described [2]. All sorted naïve B cells were CD3^−^CD19^+^IgD^+^CD27^−^ and additionally had one of the following marker profiles: T1/2 (CD10^+^MTG^+^), T3a (CD10^−^MTG^+^CD21^+^CD45RB^−^CD73^−^), T3b (CD10^−^MTG^+^CD21^+^CD45RB^−^CD73^+^), or FO (CD10^−^MTG^−^CD73^+^). Cells were plated as biological replicates in RPMI 1640 media supplemented with penicillin/streptomycin (100 U/ml), 2 mM L-glutamine, and 10% FBS. In a subset of experiments, recombinant human CD40L (TNFSF5; BioLegend) was added to the media. In Fig. 5a, amounts of (p)-S6 were analyzed at 24 hours in culture, in the absence or presence of CD40L treatment as indicated, using (p)-S6 Ser^235/236^ (monoclonal cupk43k, Thermo Fisher Scientific). Viability was assayed before permeabilization using LIVE/DEAD (Invitrogen) fixable stain according to the manufacturer’s instruction. Cells were fixed for 5 to 10 min at room temperature in 4% paraformaldehyde fixation buffer (BioLegend). Cells were permeabilized for 30 min at 4°C in PhosFlow Perm Buffer III (BD Biosciences). All subsequent steps were identical to extracellular staining.

In Fig. 5b, sorted T3a and T3b B cells were assayed for differentiation to FO at 24 hours in culture by flow cytometry using MitoTracker staining in conjunction with the following primary antibody panel: CD38 (HIT2, BD Biosciences), CD24 (ML5, BioLegend), CD10 (HI10a, BioLegend), CD45RB (MEM-55, Thermo Fisher), and CD21 (B-ly4, BD Biosciences) and CD73 (ADA, BD Biosciences). T1/2 did not significantly differentiate to FO B cells and are thus not presented in Fig. 5b.

### Bulk RNA-seq sample collection and data analysis

Total RNA was isolated from flow cytometry–sorted T1/2, T3a/b, and FO cells from three healthy individuals using the RNeasy plus Micro Kit (Qiagen). RNA-seq libraries were prepared as previously described [47]. Briefly, whole transcriptome amplification and tagmentation-based library preparation were performed using the SMART-Seq2 protocol, followed by 35–base pair (bp) paired-end sequencing on a NextSeq 500 instrument (Illumina). Five million to 10 million reads were obtained from each sample and reads were analyzed as detailed below. The data discussed in this publication were deposited in the National Center for Biotechnology Information’s Gene Expression Omnibus (GEO) with accessibility through GEO series accession no. GSE138729. Pairwise differential gene expression analysis was performed using DESeq-2 between transitional populations and follicular B cell samples [48]; p-values were generated using a Wald test statistic and adjusted using the Benjamini-Hochberg method for controlling the false discovery rate. Pathway enrichments from differentially expressed genes were performed using the Ingenuity Pathway Analysis (IPA) platform, and significant pathway overlap was determined using a right-tailed Fisher’s exact test and Benjamini-Hochberg false discovery rate correction.

### scRNA-seq with Seq-well and Computational RNA-seq pipelines

We used the Seq-Well S^3^ platform for massively parallel scRNA-seq to capture transcriptomes of single cells on barcoded mRNA capture beads. A comprehensive description of the implementation of this platform is available [49]. In brief, ∼ 15,000 IgD^+^CD27^−^ previously frozen B cells from patients with CTLA4 deficiency or healthy controls were sorted as described above and then loaded onto single arrays containing barcoded mRNA capture beads (ChemGenes). The arrays were sealed with a polycarbonate membrane (pore size of 0.01 μm), cells were lysed, transcript was hybridized, and the barcoded mRNA capture beads were recovered and pooled for all subsequent steps. Reverse transcription was performed using Maxima H Minus Reverse Transcriptase (Thermo Fisher Scientific EP0753). Exonuclease I treatment (NEB M0293 L) was used to remove excess primers. Second strand synthesis was performed using a primer of eight random bases to create complementary cDNA strands with SMART handles for PCR amplification. Whole transcriptome amplification was carried out using KAPA HiFi PCR Mastermix (Kapa Biosystems KK2602) with 2000 beads per 50-μl reaction volume. Libraries were then pooled in sets of eight (totaling 16,000 beads) and purified using Agencourt AMPure XP beads (Beckman Coulter, A63881) by a 0.6× solid phase reversible immobilization (SPRI) followed by a 0.8× SPRI and quantified using Qubit hsDNA Assay (Thermo Fisher Scientific Q32854). The quality of whole transcriptome amplification (WTA) product was assessed using the Agilent High Sensitivity D5000 Screen Tape System (Agilent Genomics) with an expected peak >800 bp tailing off to beyond 3000 bp and a small/nonexistent primer peak, indicating a successful preparation. Libraries were constructed using the Nextera XT DNA tagmentation method (Illumina FC-131–1096) on a total of 750 pg of pooled cDNA library from 16,000 recovered beads using index primers with format as previously described [49]. Tagmented and amplified sequences were purified at a 0.8× SPRI ratio yielding library sizes with an average distribution of 500 to 750 bp in length as determined using the Agilent High Sensitivity D1000 Screen Tape System (Agilent Genomics). Two arrays were sequenced per sequencing run with an Illumina 75 Cycle NextSeq500/550v2 kit (Illumina FC-404–2005) at a final concentration of 2.4 pM. The read structure was paired end with Read 1 starting from a custom Read 1 primer containing 20 bases with a 12-bp cell barcode and 8-bp unique molecular identifier (UMI) and Read 2 containing 50 bases of transcript sequence.

### Alignment & Pre-processing of scRNA-seq data

Computational pipelines and analysis read alignment were performed as previously described [49]. Briefly, for each NextSeq sequencing run, raw sequencing data were converted to demultiplexed FastQ files using bcl2fastq2 based on Nextera N700 indices corresponding to individual samples/arrays. Reads were then aligned to hg19 genome using the Galaxy portal maintained by the Broad Institute for Drop-seq Alignment using standard settings. Individual reads were tagged according to the 12-bp barcode sequenced and the 8-bp UMI contained in Read 1 of each fragment. After alignment, reads were binned onto 12-bp cell barcodes and collapsed by their 8-bp UMI. The data discussed in this publication were deposited in the National Center for Biotechnology Information’s Gene Expression Omnibus (GEO) with accessibility through GEO series accession no **GSE188449**, and on the Broad Institute Single Cell Portal under the accession number **SCP1648.**

### Single-cell RNA-sequencing clustering analysis for characterizing IgD^+^CD27^−^ naïve B cell subpopulations

Cells from each donor with greater than 500 genes or 800 UMIs were input into Seurat v3 (https://github.com/satijalab/seurat) for normalization and further analysis; this resulted in a dataset of 2,617 naïve B cell transcriptomes across three healthy donors and 2,401 naïve B cell transcriptomes across four CTLA4-deficient individuals. To avoid clustering by sequencing batch or by sample quality, variable features were carefully selected in two steps: first, 3000 variable genes were identified for each sample using Seurat’s variance-stabilizing transformation (vst) function, and the top 1500 median variable genes across all samples, excluding genes encoding ribosomal proteins, mitochondrial genes, and variable immunoglobulin light chain genes, were selected as variable features for the merged dataset. Second, to increase clustering resolution of the transitional B cell subpopulation, we identified a module of genes whose raw counts were significantly correlated to *SOX4* and *CD38* gene expression (3 standard deviations above median Pearson correlation) and significantly co-correlated with each other (Ward distance clustering of Pearson correlation values, 2 standard deviations above the median). Genes in this module were added to the sample-by-sample derived variable genes list, if they were not already included unbiasedly. scRNA-seq counts were then scaled over this curated variable features gene list for downstream clustering analysis.

We then performed principal component analysis (PCA) and applied a JackStraw test to determine a significant number of PCs (n=6) to use for both uniform manifold approximation and projection (UMAP) and SNN (shared nearest neighbor) clustering, removing one PC that exclusively defined low quality samples to avoid introducing technically-driven batch effect into the clustering. Transitional B and Age-associated B cell clusters were separated from follicular B cells using Louvain clustering and UMAP visualization with a resolution of 0.4 and k=30.

Normalized counts were used for marker gene visualization, and adjusted *P* values for significant differentially expressed genes were generated using a non-parametric Wilcoxon rank sum test and Bonferroni multiple hypothesis correction. Finally, module scoring was performed using Seurat’s AddModuleScore function to construct a mean score of supplied genes subtracting a background score constructed from a random selection of genes in bins of average expression across all cells. Signatures for oxidative phosphorylation and mTORC1 signaling were obtained from the msigDB database (HALLMARK_OXIDATIVE_PHOSPHORYLATION, HALLMARK_MTORC1_SIGNALING), and statistical significance was calculated using a pairwise non-parametric Wilcoxon rank sum test and Benjamini Hochberg false discovery rate correction.

### Multi-color immunofluorescence staining

Tissue samples were fixed in formalin, embedded in paraffin, and sectioned. These specimens were incubated with the following antibodies: anti-CD19 (1:200; SKU310; Biocare Medical), anti-IgD (1:2000; AA093; DAKO), anti-CD27 (1:1000; clone: ab131254; Sigma Prestige), anti-CD40L (1:100; ab2391; Abcam). The samples were mounted with ProLong Diamond Antifade mountant containing DAPI (Invitrogen).

### Microscopy and Quantitative Image Analysis

Images of the tissue specimens were acquired using the TissueFAXS platform (TissueGnostics). For quantitative analysis, the entire area of the tissue was acquired as a digital grayscale image in five channels with filter settings for FITC, Cy3, Cy5 and AF750 in addition to DAPI. Cells of a given phenotype were identified and quantitated using the TissueQuest software (TissueGnostics), with cut-off values determined relative to the positive controls. This microscopy-based multicolor tissue cytometry software permits multicolor analysis of single cells within tissue sections similar to flow cytometry. In addition, multispectral images (seven-colors staining) were unmixed using spectral libraries built from images of single stained tissues for each reagent using the StrataQuest (TissueGnostics) software. StrataQuest software was also used to quantify cell-to-cell contact. In the StrataQuest cell-to-cell contact application, masks of the nuclei based on DAPI staining establish the inner boundary of the cytoplasm and the software ‘‘looks’’ outward toward the plasma membrane boundary. Overlap of at least 3 pixels of adjacent cell markers is required to establish a ‘‘contact’’ criterion. The number of fields examined depends on the size of the sample, but is at least 100 mm^2^. Although the software has been developed and validated more recently, the principle of the method and the algorithms used have been described in detail elsewhere (Ecker and Steiner, 2004).

### Statistical methods & analysis

Statistical analysis was conducted using GraphPad Prism version 9.00 and statistical tests chosen according to the specific context. Student’s t-test was used to compare two groups of parametric variables while two-tailed Mann-Whitney U test was used to calculate *P* values for continuous, non-parametric variables. ANOVA with post-hoc Tukey test was used for multiple comparisons with paired Wilcoxon rank sum test for pair-wise comparisons. Correlation by nonparametric Spearman’s rank with correlation coefficient (r) was used for correlation of nonparametric variables. *P* value inferior to 0.05 was considered as significant.

## Supplementary Materials

**Fig. S1:** Flow Cytometry Gating Strategies

**Fig. S2:** CTLA4 levels in human transitional and follicular B cells

**Fig. S3:** Characterization of T cell subsets in CTLA4 deficiency

**Fig. S4:** T cell phenotype after abatacept therapy

**Fig. S5:** CCR7 is upregulated at the T3a stage in human B cells

**Fig. S6:** CD86 levels in naïve (IgD^+^CD27^−^) human B cells

**Fig. S7:** CD40L+ T cells interact with naïve B cells in lymphoid tissue of patients with congenital T cell hyperactivation

**Table S1:** Clinical characteristics of patients with CTLA4 deficiency

**Table S2:** B cell panel for the analysis of PBMCs from patients with CTLA4 deficiency

**Table S3**: Extra-cellular T cell panel for the analysis of PBMCs from patients with CTLA4 deficiency

**Table S4**: Combined intra-/extra-cellular T cell panel for the analysis of PBMCs from patients with CTLA4 deficiency

## Acknowledgments

We thank Drs. Bodo Grimbacher and David Sansom for kindly providing the CHO cells stably overexpressing CD80-GF [3] and the tissue processing core at the Ragon Institute for the processing and biobanking of the CTLA4-deficient patient samples. **Funding** was provided by the American Society of Hematology Research Training Award for Fellows (KH), NIH K23AI163350 (SB), NIH U19 AI110495 and the Ragon Insitute (SP), and NIH T32-HL116275 and Bristol Myers Squibb (JRF). The content is solely the responsibility of the authors and does not necessarily represent the official views of the National Institutes of Health.

## Author contributions

Conceptualization: HAC, JRF, SP. Patient sample contribution: SB, EWC, JW, AS. Methodology: JRF, KH, HAC, AB, TL, CT, MR, GY, VMRC, KP, NK, MG, Funding acquisition: KH, JRF, SP. Supervision: AKS, JRF, SP. Writing – original draft: HAC, KH, JRF, Writing – review & editing: all authors.

## Data and materials availability

All data needed to evaluate the conclusions in the paper are present in the main paper or the supplementary materials.

