## Supplemental Materials for "Congenital T cell activation impairs transitional to follicular B cell maturation in humans"

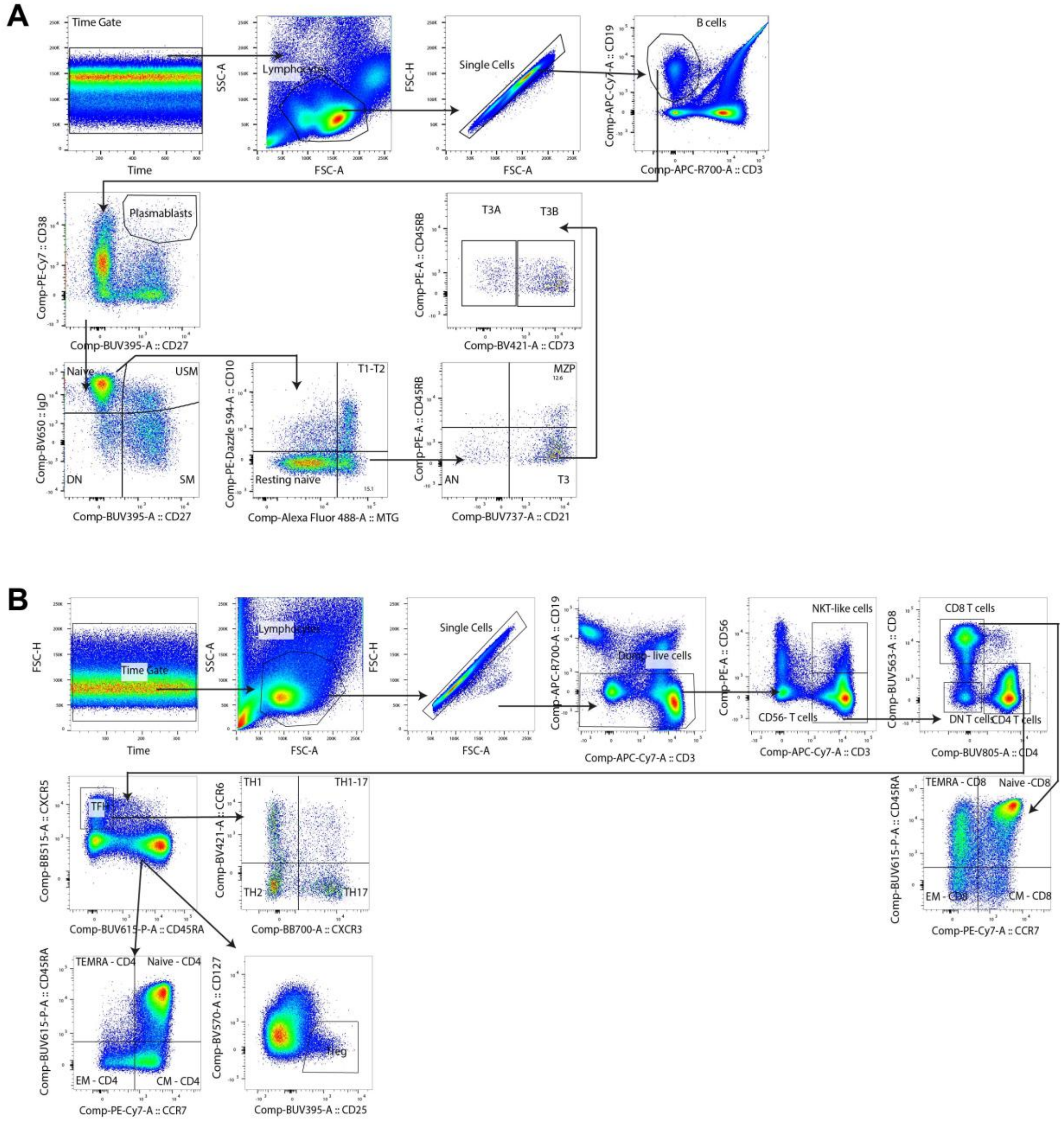


**Supplementary Fig. 1. Flow Cytometry Gating Strategies.** Flow cytometry gating strategies for (A) B cells and (B) T cells. (A) After gating on stable event and lymphocytes, doublets and dead cells were removed. CD27$^{++}$/CD38$^{++}$ plasmablasts where gated out first, then B cells were subset as Naïve (IgD$^{+}$CD27$^{-}$), unswitched memory (USM; IgD$^{+}$CD27$^{+}$), double negative (DN; IgD$^{-}$CD27$^{-}$) and switched memory (SM; IgD$^{-}$CD27$^{+}$). Naïve B cells were further divided to follicular (FO), transitional (T1-3b), activated naïve (AN), and marginal zone precursor (MZP) using the markers MTG, CD10, CD73, CD21 and CD45RB as shown. (B) After gating on stable event and lymphocytes, doublets and dead cells were removed. From CD3$^{+}$ T cells, NKT-like cells were first gated out using the marker CD56$^{+}$. Next, CD4$^{+}$ and CD8$^{+}$ T cells were subset, excluding double negative ($CD4^{-}CD8^{-}$) T cells. $CD4^{+}$ T follicular helper (TFH) cells were identified using $CXCR5^{+}$ as a marker and TFH were further subdivided into TH1, TH2, and TH17 using CXCR3 and CCR6 as markers. The remaining $CD4^{+}$ T cells and $CD8^{+}$ T cells were gated using CD45RA and CCR7 to separate Naïve ($CD45RA^{+}CCR7^{+}$), central memory (CM; $CD45RA^{-}CCR7^{+}$), effector memory (EM; $CD45RA^{-}CCR7^{-}$) and effector memory $CD45RA^{+}$ (TEMRA; $CD45RA^{+}CCR7^{-}$). T regulatory (Treg) cells were gated from $CD4^{+}CXCR5^{-}$ T cells as $CD127^{lo}CD25^{hi}$ cells.

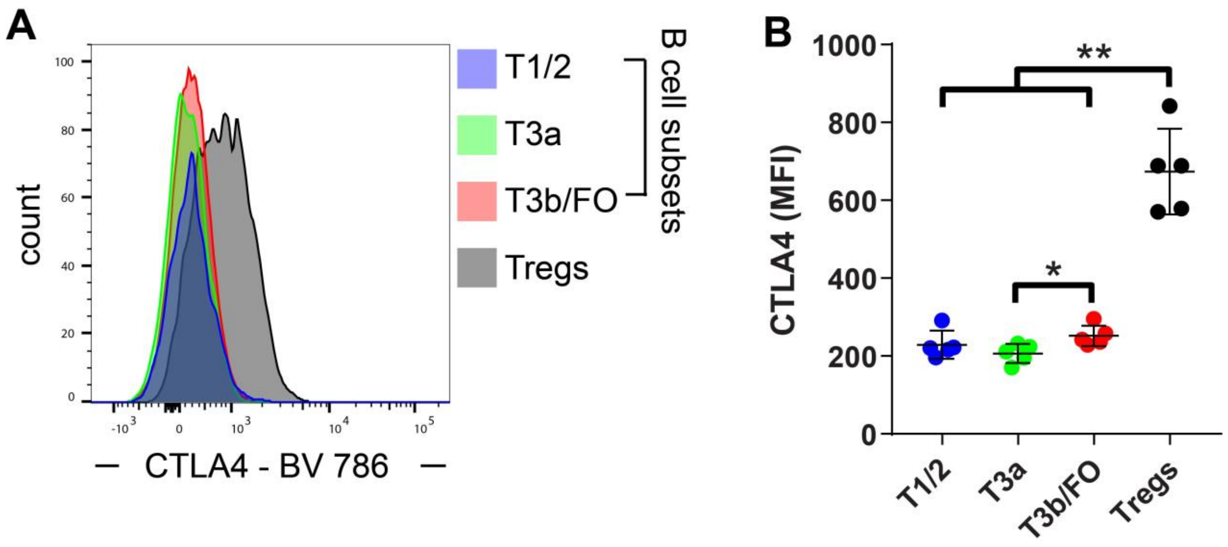


**Supplementary Fig. 2. CTLA4 levels in human transitional and follicular B cells.** Flow cytometry analysis of CTLA4 levels by intracellular flow cytometry in human transitional (T1-3b) and follicular (FO) B cell subsets and T regulatory cells (Tregs) in healthy controls (n=5). Data shown as histograms (A) as well as quantification and analysis (B). Symbols represent unique individuals; bars represent means (+/- SD) of all data. *$P < 0.05$; **$P < 0.01$ by one-way ANOVA.

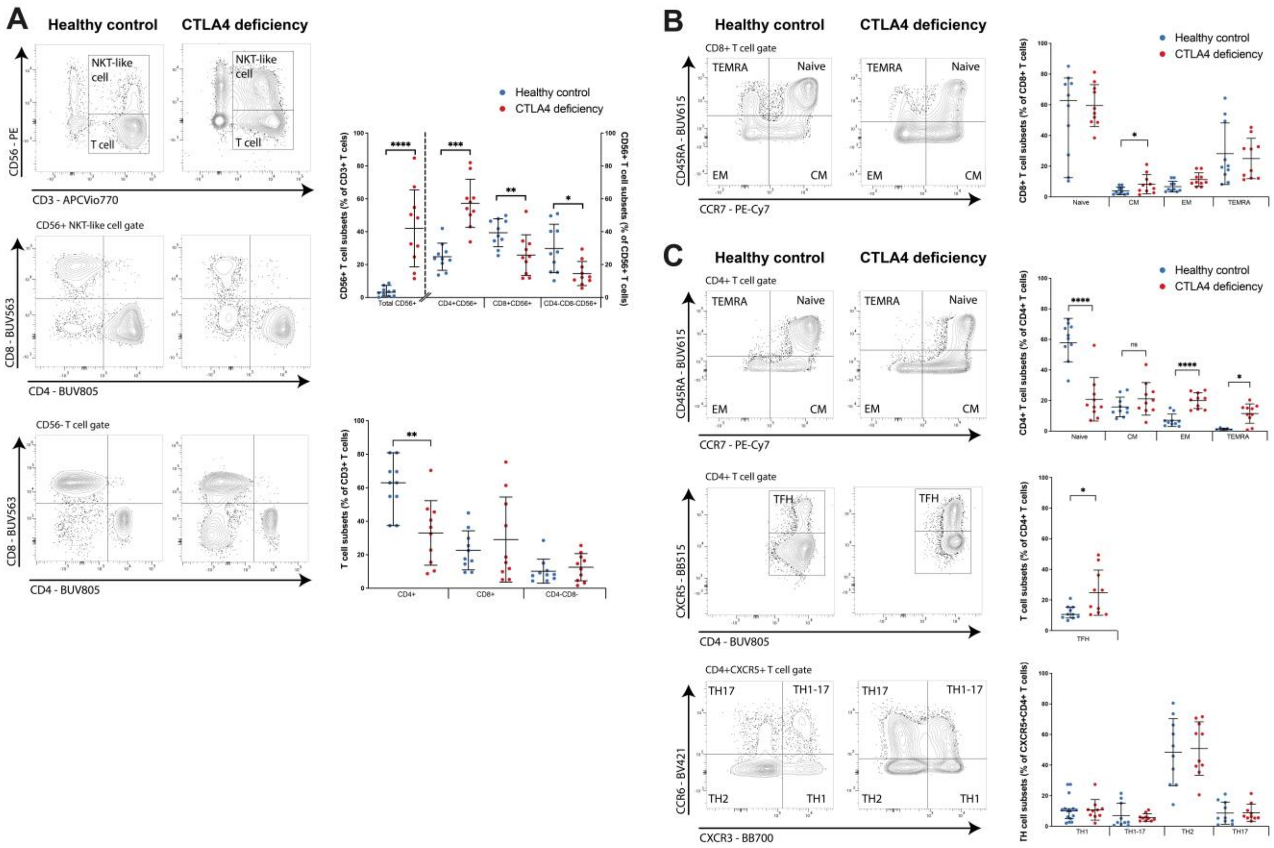


**Supplementary Fig. 3. Characterization of T cell subsets in CTLA4 deficiency.** Flow cytometric analysis of T cell populations in peripheral blood in 10 CTLA4-deficient patients as compared to 10 healthy controls. (A) Analysis and quantification of CD56$^+$ NKT-like cells and CD56$^-$ T cells. (B) Analysis and quantification of CD8$^+$ T cell gate. (C) Analysis and quantification of CD4$^+$ T cell gate. Contour plots are representative of 10 healthy controls and CTLA4-deficient patients, respectively. Symbols represent unique individuals; bars represent means (+/-) SD of all data. *$P$ < 0.05; **$P$ < 0.01; ***$P$ < 0.001; ****$P$ < 0.0001 by two-tailed Mann-Whitney U test.

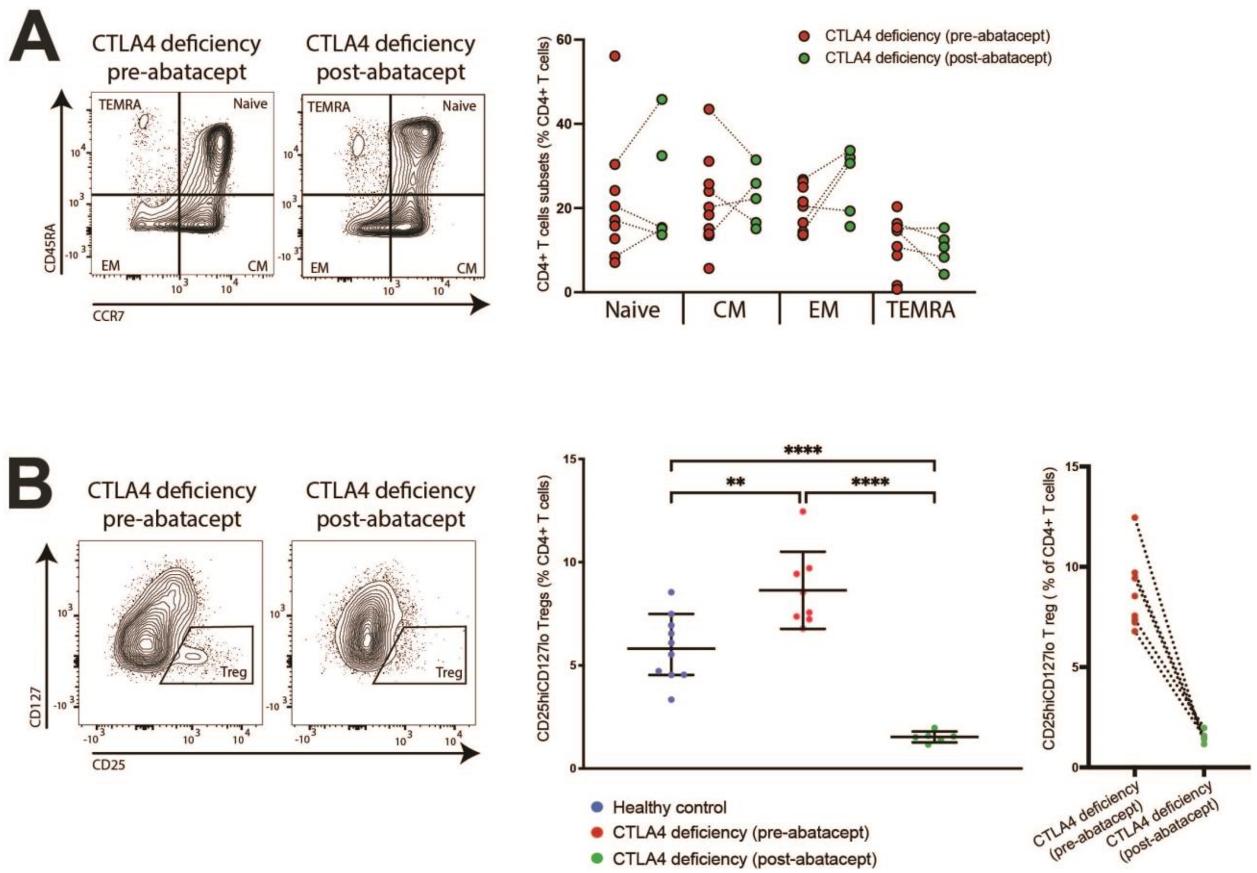


**Supplementary Fig. 4. T cell phenotype after abatacept therapy.** Flow cytometric analysis of T cell populations in CTLA4-deficient patients pre- (n=8) and post-treatment (n=6) with abatacept therapy. (A) Analysis and quantification of CD4+ T cell subsets in peripheral blood (naïve, central memory (CM), effector memory (EM), and T effector memory RA (TEMRA)). (B) Analysis and quantification of $CD127^{lo}CD25^{hi}$ Treg cell gate in peripheral blood. Contour plots are representative of 8 CTLA4-deficient patients pre-abatacept treatment as compared to 6 CTLA4-deficient patients post-abatacept treatment, respectively. Symbols represent unique individuals; bars represent means (+/-) SD of all data. Dotted lines indicate a paired pre- to post-treated individual patient. *$P < 0.05$; **$P < 0.01$; ***$P < 0.001$; ****$P < 0.0001$ by ANOVA.

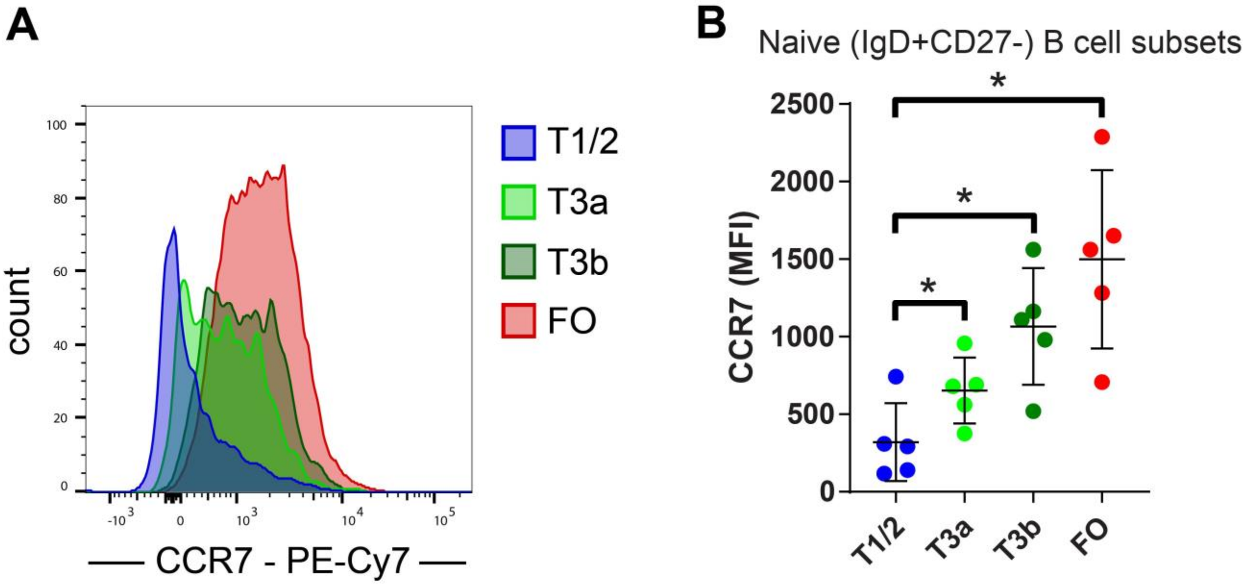


**Supplementary Fig. 5. CCR7 is upregulated at the T3a stage in human B cells.** Flow cytometric analysis of CCR7 surface levels in human transitional (T1-3b) and follicular (FO) B cell subsets in healthy controls (n=5). Data shown as histograms (A) as well as quantification and analysis (B). Symbols represent unique individuals; bars represent means (+/- SD) of all data. **P*<0.05 by one-way ANOVA.

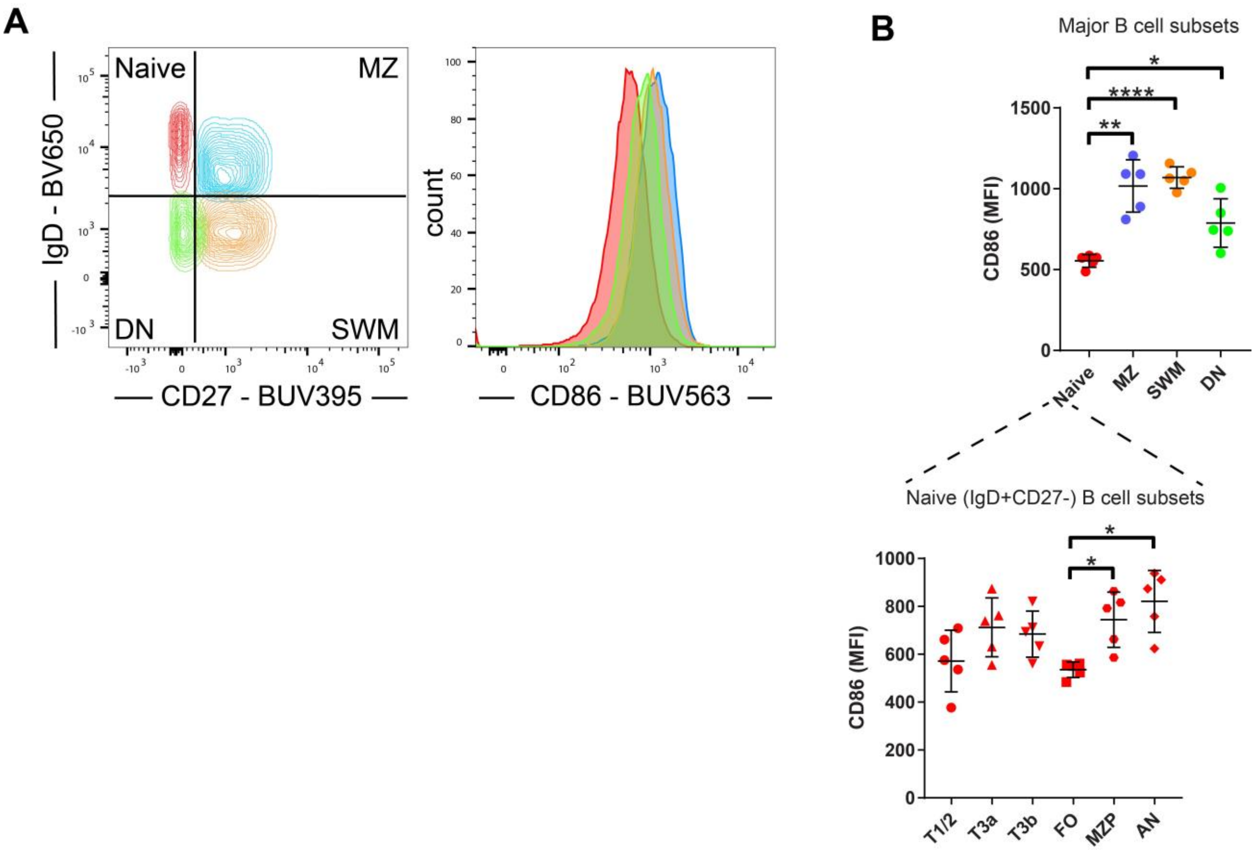


**Supplementary Fig. 6. CD86 levels in naïve ($IgD^{+}CD27^{-}$) human B cells.** Flow cytometric analysis of CD86 surface levels in human B cell subsets (naïve, marginal zone (MZ), double negative (DN), switched-memory (SWM), transitional (T1-3b), follicular (FO), marginal zone precursor (MZP), and activated naïve (AN)) in healthy controls (n=5). Data shown as contour plot and histogram (A) as well as quantification and analysis (B). Symbols represent unique individuals; bars represent means (+/- SD) of all data. *$P < 0.05$; **$P < 0.01$; ***$P <$ 0.001; ****$P < 0.0001$by ANOVA.

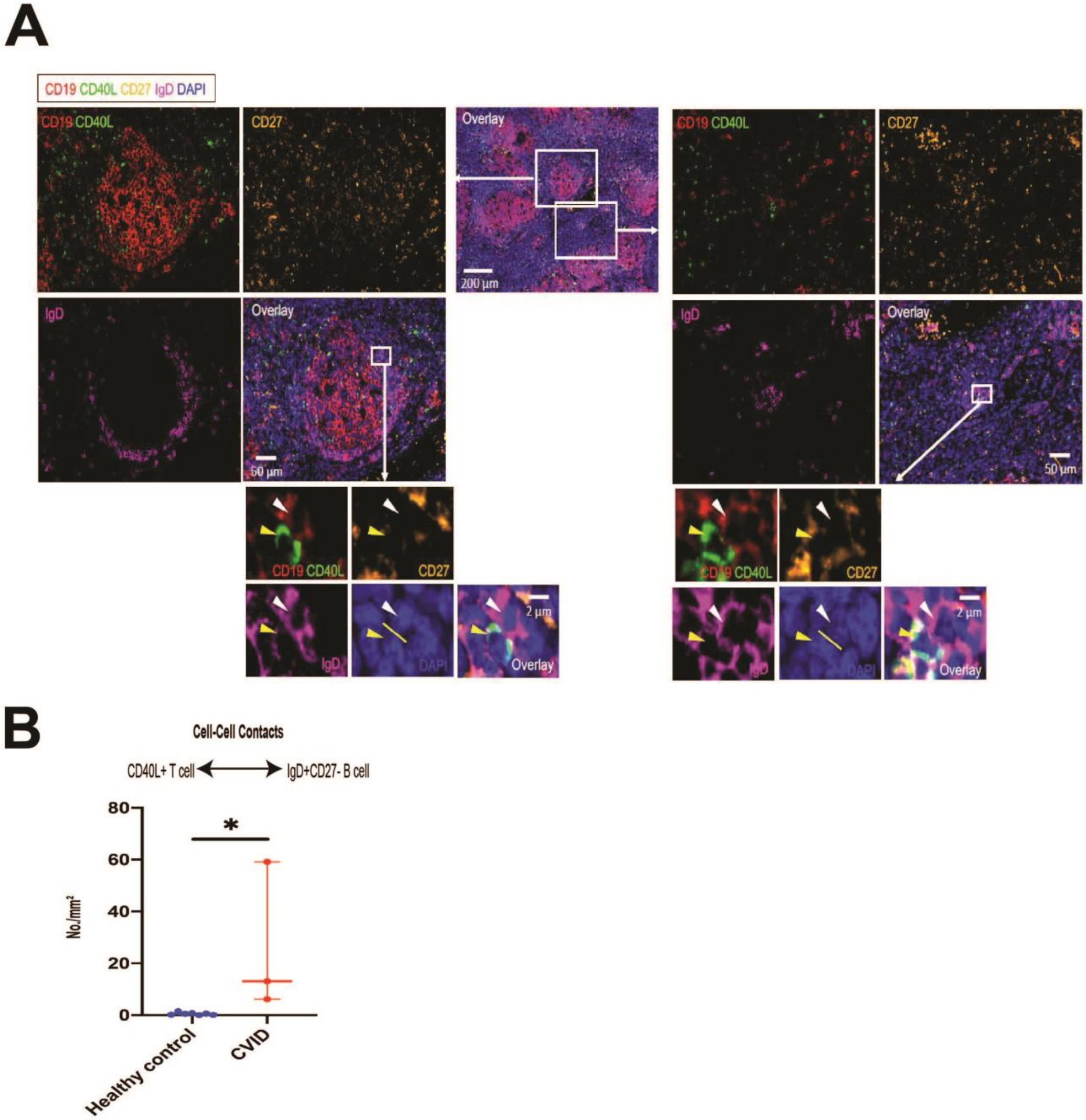


**Supplementary Fig. 7: CD40L+ T cells interact with naïve B cells in lymphoid tissue of patients with congenital T cell hyperactivation.** (A) Representative multi-color immunofluorescence images of CD19 (red), CD40L (green), CD27 (orange), IgD (purple) and DAPI (blue) staining of a lymph node from a patient with CTLA4 deficiency presenting clinically as common variable immunodeficiency (CVID). The yellow bar is placed between the nucleus of a CD40L expressing cell (below left) and the nucleus of a naïve $CD19^{+}IgD^{+}CD27^{-}$ B cell (above right). Yellow arrows delineate T cells in selected T-B conjugates while B cells are delineated by white arrows. (B) Absolute numbers of cell-cell contacts between CD40L+ T cells and $IgD^{+}CD27^{-}$ B cells in healthy control tonsils (blue) (n=7) and lymph nodes of patients with congenital T cell activation, due to activated PI3K delta syndrome (APDS) or CTLA4 deficiency (red) (n = 3) are shown. Quantified data are means (+/- SD) of all data. *$P < 0.05$by Wilcoxon rank sum test (B).

| **Family** | 1 | 1 | 1 | 2 | 3 | 3 | 3 | 4 | 4 | 4 | 5 | 5 | 5 |
|---|---|---|---|---|---|---|---|---|---|---|---|---|---|
| **Member** | proband | sister | brother | proband | proband | son 1 | son 2 | proband | brother | sister | cousin | Son of cousin | proband |
| **CTLA4 mutation** | 3' UTR mutation (homo.) | 3' UTR mutation (homo.) | 3' UTR mutation (het.) | deletion 2q33.1 - q33.3 (het.) | c.410C>T, p.P137L (het.) | c.410C>T, p.P137L (het.) | c.410C>T, p.P137L (het.) | c.81dupT, p.L28SfsX402 (het.) | c.81dupT, p.L28SfsX402 (het.) | c.81dupT, p.L28SfsX402 (het.) | c.173G>C, p.C58S (het.) | c.173G>C, p.C58S (het.) | c.173G>C, p.C58S (het.) |
| **Other Genetic Variants** | None by WES | None by WES | None by WES | | None by WES | None by WES | None by WES | *TNFRSF13B* (p.A181E) *NOD2* (p.L1007PfaX2), ADA2 (deletion exon 7) | ADA2 (deletion exon 7) | *TNFRSF13B* (p.A181E) *NOD2* (p.L1007PfaX2) | Full PID panel still needs to be done. This patient was tested via family-variant testing program – her cousin is proband | CHD7 | Negative Invitae ALPS gene panel |
| **Age at B cell immunophenotyping** | 23 years | 25 years | 31 years | 25 years | 45 years | 20 years | 17 years | 13 years | 8 years | 19 years | 49 years | 13 years | 51 years |
| **Gender** | Male | Female | Male | Male | Female | Male | Male | Male | Male | Female | Female | Male | Female |
| **Lymphoproliferation/ Autoimmunity** | GLILD, NRH, splenomegaly, LAD, AI enteropathy, DMI, Coombs positive AIHA, ITP | Seronegative inflammatory arthritis | (none) | AI enteropathy, inflammatory arthritis NOS | Hashimoto's thyroiditis, seronegative inflammatory arthritis | (none) | LAD, IBD | Colitis, inguinal lymphadenopathy, splenomegaly | (none) | AIHA, pancreatitis, colitis on CT (not yet confirmed on biopsy) | IBD, inflammatory arthritis, ILD, ITP, prominent mediastinal/hilar/axillary/retroperitoneal lymphadenopathy; bilateral pulmonary nodules,migratory airspace consolidation, thick mucoid secretions on bronchoscopy | asthma | ITP, WAIHA, Hypogammaglobulinemia, GL-ILD, Mediastinal and extensive hilar lymphadenopathy, AI enteropathy |
| **Bronchiectasis** | Yes | No | n/a | Yes | No | n/a | No | No | No | No | No | No | No |
| **ALC (1000-4800 cells/μL)** | | | n/a | | | n/a | | 1166 | 2122 | 756 | 955 | 3172 | 1060 |
| **$CD3^+$ (690-2540 cells/μL)** | 275 | 1077 | n/a | 956 | 255 | n/a | 955 | 791 (910-2200) | 1457 | 583 | 762 | 2208 | n/a |
| **$CD4^+$ (419-1590 cells/μL)** | 160 | 642 | n/a | 608 | 139 | n/a | 434 | 357 (520-1440) | 940 | 195 | 598 | 1324 | n/a |
| **$CD4^+CD45RA^+$ (% $CD4^+$)** | 1.7 | | n/a | 26.5 | 1.1 | n/a | 24.4 | n/a | n/a | 14 | 6 | 34 | n/a |

| | | | | | | | | | | | | | |
|---|---|---|---|---|---|---|---|---|---|---|---|---|---|
| **$CD4^{+}CD45RO^{+}$ (% $CD4^{+}$)** | 97.1 | | n/a | 62.8 | 87.4 | n/a | 67.5 | n/a | n/a | n/a | n/a | n/a | n/a |
| **$CD8^{+}$ (190-1140 cells/μL))** | 112 | 400 | n/a | 194 | 107 | n/a | 366 | 378 (300-900) | 454 | 375 | 119 | 745 | n/a |
| **$CD8^{+}CD45RA^{+}$ (% CD8+)** | 22.4 | | n/a | 53.2 | 11.6 | n/a | 42.5 | n/a | n/a | 16 | 5 | 24 | n/a |
| **$CD8^{+}CD45RO^{+}$ (% $CD8^{+}$)** | 68.8 | | n/a | 30.5 | 60.6 | n/a | 54.1 | n/a | n/a | n/a | n/a | n/a | n/a |
| **$CD19^{+}$ (90-660 cells/μL)** | 14 | 86 | n/a | 539 | 33 | n/a | 484 | 272 (200-820) | 404 | 119 | 114 | 727 | n/a |
| **$CD3^{-}CD16/56^{+}$ (90-590 cells/μL)** | 55 | 122 | n/a | 140 | 128 | n/a | 208 | 64 (70-500) | 206 | 54 | 78 | 214 | n/a |
| **$CD27^{+}$ (% $CD19^{+}$)** | 4 | 5 | n/a | 16.6 | 22.1 | n/a | 10.3 | 24 | 19 | 16 | 12 | 23 | n/a |
| **$CD27^{+}IgD/M^{-}$ (% $CD19^{+}$)** | 0.4 | <1 | n/a | 1.8 | 4 | n/a | 1.7 | n/a | n/a | n/a | n/a | n/a | n/a |
| **IgG (614-1295 mg/dL)** | 422 (L) | 696 | n/a | 119->679 (control of protein wasting) | 567 | n/a | 648 | 748 (893-1823) | 662 | 267 | 512 | 979 | 544 |
| **IgA (69-309 mg/dL)** | 7 | 95 | n/a | 5 | 38 | n/a | 52 | 59 (70-432) | 105 | 11 | 11 | 84 | 167 |
| **IgM (53-334 mg/mL)** | 5 | 358 (H) | n/a | 214 | 66 | n/a | 71 | 15 (52-367) | 14 | 25 | 52 | 62 | 143 |
| **Pneumococcal titers (≥ 1.3 μg/mL)** | | 2 of 23 serotypes (8.7 %) | n/a | 70% | 82.6 % | n/a | | 0 of 23 $\rightarrow$ 0 of 23 | 0 of 14$\rightarrow$ 8 of 14 In 2013 | 0 of 23$\rightarrow$4 of 23 | 2 of 23 | 4 of 23 $\rightarrow$21/23 | |
| **Proliferation to PHA** | Decreased | Normal | n/a | | | n/a | | Normal Date 2/20/18 | n/a | n/a | Normal (high) | Normal | |
| **Proliferation to PWM** | Insufficient | Insufficient | n/a | | | n/a | | Normal | n/a | n/a | Normal | Normal | |
| **Proliferation to Candida** | Insufficient | Normal | n/a | | | n/a | | Normal | n/a | n/a | Normal | Normal | |
| **Proliferation to Tetanus** | Insufficient | Normal | n/a | | Normal | n/a | | Insufficient | n/a | n/a | Normal | Normal | |
| **Proliferation to aCD3/IL2** | Normal | | n/a | | | n/a | | n/a | n/a | n/a | n/a | n/a | |

| | | | | | | | | | | | | | |
|---|---|---|---|---|---|---|---|---|---|---|---|---|---|
| **Treatment prior to B cell phenotyping** | IVIG | IVIG (Tofacitinib starting in 2016) | None | None | Prednisone | None | None | IVIG | n/a | None (sample rejected for hemolysis) | n/a | n/a | Dexamethasone, Prednisone, IVIG, Rituximab, Romiplostim |
| **Abatacept Treatment** | Started on 3/8/16 (125 mg SC weekly) | None | None | Started on 5/20/16 (125 mg SC weekly) | 9/25/18 (1000 mg IV q 2 weeks) | None | Started on 2/1/18 (1350 mg IV monthly) | Started on 07/11/2019 | None | Started on 06/27/2019 on emergency basis for AIHA | Started 1/16/2020 | None | Started on 11/1/2019 |
| **B cell Panel** | Yes | Yes | Yes | Yes (only post-abatacept) | Yes | Yes | Yes | Yes | Yes | Yes (only post-abatacept) | Yes | Yes | Yes |
| **Extracellular T cell Panel** | Yes | Yes | Yes | Yes (only post-abatacept) | Yes | Yes | Yes | Yes | No | No | Yes | Yes | Yes |
| **Intracellular T cell Panel** | No | No | No | Yes (only post-abatacept) | Yes | Yes | Yes | No | No | No | Yes | Yes | Yes |
| **Transendocytosis** | No | Yes | Yes | Yes | Yes | Yes | Yes | No | No | Yes | Yes | Yes | Yes |

Normal reference ranges from the Massachusetts General Hospital shown where applicable. Pneumococcal titers shown as ratio of positive over tested serotypes. Autoimmune enteropathy (AIE); autoimmune hemolytic anemia (AIHA); immune thrombocytopenia (ITP); intravenous immunoglobulin (IVIG); female (F); granulomatous-lymphointerstitial lung disease (GLILD); human papilloma virus (HPV); lymphadenopathy (LAD); male (M); not available (n/a); subcutaneous immunoglobulin (SCIG); varicella zoster virus (VZV).

**Supplemental Table 1.** Clinical characteristics of patients with CTLA4 deficiency.

| Antigen | Fluorophore | Clone | Supplier | Dilution | Staining Condition |
|---|---|---|---|---|---|
| CD27 | BUV395 | L128 | BD Biosciences | 1:40 | EC – 4ºC |
| IgM | BUV496 | G20-127 | BD Biosciences | 1:80 | EC – 4ºC |
| IgG | BUV563 | G18-145 | BD Biosciences | 1:100 | EC – 4ºC |
| CD21 | BUV737 | B-ly4 | BD Biosciences | 1:20 | EC – 4ºC |
| CD20 | BUV805 | 2H7 | BD Biosciences | 1:60 | EC – 4ºC |
| CD73 | BV421 | ADA | BD Biosciences | 1:40 | EC – 4ºC |
| CXCR5 | BV605 | J252D4 | BioLegend | 1:30 | EC – 4ºC |
| IgD | BV650 | IA6-2 | BD Biosciences | 1:80 | EC – 4ºC |

| | | | | | |
|---|---|---|---|---|---|
| CD11c | BV711 | B-ly6 | BD Biosciences | 1:20 | EC – 4$^{o}$C |
| MitoTracker Green (MTG) | AF488 | - | ThermoFisher | 100-nM | Stain/Efflux – 37$^{o}$C |
| CD24 | BB700 | ML5 | BioLegend | 1:60 | EC – 4$^{o}$C |
| CD45RB | PE | MEM-55 | ThermoFisher | 1:20 | EC – 4$^{o}$C |
| CD10 | PE-Dazzle594 | HI10a | BioLegend | 1:20 | EC – 4$^{o}$C |
| CD38s | PE-Cy7 | HIT2 | BioLegend | 1:30 | EC – 4$^{o}$C |
| CD3 | APC-R700 | UCHT1 | BD Biosciences | 1:30 | EC – 4$^{o}$C |
| CD19 | APC-Cy7 | SJ25C1 | BioLegend | 1:30 | EC – 4$^{o}$C |

**Supplemental Table 2.** B cell panel for the analysis of PBMCs from patients with CTLA4 deficiency.

| **Antigen** | **Fluorophore** | **Clone** | **Supplier** | **Dilution** | **Staining Condition** |
|---|---|---|---|---|---|
| CD25 | BUV395 | 2A3 | BD Biosciences | 1:20 | EC – 4$^o$C |
| CD57 | BUV496 | NK-1 | BD Biosciences | 1:200 | EC – 4$^o$C |
| CD45RA | BUV615 | HI100 | BD Biosciences | 1:300 | EC – 4$^o$C |
| CD28 | BUV737 | CD28.2 | BD Biosciences | 1:20 | EC – 4$^o$C |
| CD4 | BUV805 | SK3 | BD Biosciences | 1:40 | EC – 4$^o$C |
| CCR6 | BV421 | G034E3 | BioLegend | 1:50 | EC – 37$^o$C |
| TCR-dg | BV480 | B1 | BD Biosciences | 1:40 | EC – 4$^o$C |
| CD127 | BV570 | A019D5 | BioLegend | 1:20 | EC – 4$^o$C |
| CD38 | BV650 | HIT2 | BD Biosciences | 1:20 | EC – 4$^o$C |
| PD-1 | BV711 | EH12.1 | BD Biosciences | 1:20 | EC – 4$^o$C |
| CD8 | BV786 | RPA-T8 | BD Biosciences | 1:400 | EC – 4$^o$C |
| CXCR5 | BB515 | RF8B2 | BD Biosciences | 1:50 | EC – 37$^o$C |

| | | | | | |
|---|---|---|---|---|---|
| CCR4 | BB630 | 1G1 | BD Biosciences | 1:20 | EC – 37$^{o}$C |
| CXCR3 | BB700 | 1C6/CXCR3 | BD Biosciences | 1:10 | EC – 37$^{o}$C |
| CD56 | PE | MEM-188 | BioLegend | 1:20 | EC – 4$^{o}$C |
| CD27 | PE-Dazzle594 | M-T271 | BioLegend | 1:40 | EC – 4$^{o}$C |
| HLA-DR | PE-Cy5.5 | TU36 | ThermoFisher | 1:20 | EC – 4$^{o}$C |
| CCR7 | PE-Cy7 | G043H7 | BioLegend | 1:10 | EC – 37$^{o}$C |
| SLAMF7 | AF647 | 235614 | BD Biosciences | 1:10 | EC – 4$^{o}$C |
| CD19 | APC-R700 | SJ25C1 | BD Biosciences | 1:30 | EC – 4$^{o}$C |
| CD3 | APC-Vio770 | BW264/56 | Miltenyi Biotec | 1:40 | EC – 4$^{o}$C |

**Supplemental Table 3.** Extra-cellular T cell panel for the analysis of PBMCs from patients with CTLA4 deficiency.

| Antigen | Fluorophore | Clone | Supplier | Dilution | Staining Condition |
|---|---|---|---|---|---|
| CD25 | BUV395 | 2A3 | BD Biosciences | 1:20 | EC – 4$^{o}$C |
| CD8 | BUV563 | RPA-T8 | BD Biosciences | 1:400 | EC – 4$^{o}$C |
| CD45RA | BUV615 | HI100 | BD Biosciences | 1:300 | EC – 4$^{o}$C |
| CD4 | BUV805 | SK3 | BD Biosciences | 1:40 | EC – 4$^{o}$C |
| CCR6 | BV421 | G034E3 | BioLegend | 1:50 | EC – 37$^{o}$C |
| CD40L | BV510 | 24-31 | BioLegend | 1:20 | IC |
| CD127 | BV570 | A019D5 | BioLegend | 1:20 | EC – 4$^{o}$C |
| CD62L | BV650 | SK11 | BD Biosciences | 1:200 | EC – 4$^{o}$C |
| CTLA4 | BV786 | BNI3 | BD Biosciences | 1:20 | IC |
| CXCR5 | BB515 | RF8B2 | BD Biosciences | 1:50 | EC – 37$^{o}$C |
| CXCR3 | BB700 | 1C6/CXCR3 | BD Biosciences | 1:10 | EC – 37$^{o}$C |
| CD56 | PE | MEM-188 | BioLegend | 1:20 | EC – 4$^{o}$C |

| | | | | | |
|---|---|---|---|---|---|
| CCR7 | PE-Cy7 | G043H7 | BioLegend | 1:10 | EC – 37°C |
| FoxP3 | AF647 | 236A/E7 | BD Biosciences | 1:20 | IC |
| CD19 | APC-R700 | SJ25C1 | BD Biosciences | 1:30 | EC – 4°C |
| CD3 | APC-Vio770 | BW264/56 | Miltenyi Biotec | 1:40 | EC – 4°C |

**Supplemental Table 4.** Combined intra-/extra-cellular T cell panel for the analysis of PBMCs from patients with CTLA4 deficiency.
